# *Lotus japonicus* VIH2 is an inositol pyrophosphate synthase that regulates arbuscular mycorrhiza

**DOI:** 10.1101/2024.12.17.628921

**Authors:** Kiran Raj, Verena Gaugler, Mengsi Lu, Maren Schädel, Philipp Gaugler, Charlotte M. M. Grothaus, Ulrike A. Jochimsen, Guizhen Liu, Michael Harings, Henning J. Jessen, Gabriel Schaaf, Martina K. Ried-Lasi

**Affiliations:** Leibniz Institute of Plant Biochemistry, Department of Molecular Signal Processing, 06120 Halle (Saale), Germany; University of Bonn, Institute of Crop Science and Resource Conservation (INRES), 53115 Bonn, Germany; Albert-Ludwigs-University of Freiburg, Institute of Organic Chemistry and the Center for Integrative Biological Signalling Studies (CIBSS), 79104 Freiburg, Germany

## Abstract

Plant yield is often maximized by the extensive use of mineral fertilizers, which, however, has severe environmental consequences. Phosphate is particularly problematic, as it represents a globally limited resource, and its runoff and soil erosion threaten open water bodies. Many crops engage in arbuscular mycorrhizal (AM) symbiosis with nutrient-acquiring fungi, aiding in the uptake of phosphate and other mineral nutrients. However, AM colonization is strongly reduced under high soil phosphate levels. A mechanistic understanding of phosphate sensing, phosphate starvation responses, and their connection to AM remains enigmatic. Here, we show that in *Lotus japonicus*, low-abundant, energy-rich inositol pyrophosphates act as master regulators of AM, orchestrating the crosstalk between phosphate starvation responses and plant root endosymbiosis. These findings hold promise for breeding nutrient-efficient crops.

## Introduction

The macronutrient phosphorus (P) is a key determinant of crop performance and yield. Inorganic phosphate (P_i_), the P-form taken up by plants, quickly becomes immobilized in most soils, making the application of P-fertilizers critical to maintain crop productivity (*1*). However, P-fertilization comes with environmental costs as global P-deposits are limited. Furthermore, soil erosion contributes to eutrophication of open water bodies with P, a major stressor on marine ecosystems (*2*).

Plants respond to P_i_ limitations through local and systemic phosphate starvation responses (PSRs) that involve morphological, transcriptional, and metabolic changes. Local responses primarily alter root system architecture, while systemic responses regulate P_i_ homeostasis (*1*). Arabidopsis Phosphate Starvation Response Regulator1 (PHR1; (*3*)) and related transcription factors (PHR1-likes; PHLs; (*4–6*)) are key regulators of PSRs and orthologs have been identified in several plant species (*7–11*). While under P_i_ limiting conditions, PHR1 targets specific motifs in the promoters of P_i_ starvation-induced genes thus activating their expression (*3*), under P_i_ sufficiency PHR1 interacts with SYG1/Pho81/XPR1 (SPX) domain-containing proteins, which prevents the binding to its promoter core sequences (*12–17*). SPX domains are selective high-affinity receptors for inositol pyrophosphates (PP-InsPs), which mediate the interaction between SPX and PHR in PSR regulation (*18–20*).

PP-InsPs (such as InsP_7_ and InsP_8_) derive from inositol hexakisphosphate (phytate, InsP_6_), which serves as the primary P-storage molecule in seeds and, aside from ATP, likely also in the whole plant (*21*). They are energy-rich, low-abundant messengers found in all eukaryotes (*22*) and act as proxies for plant P_i_ status (*20*, *23*, *24*). In Arabidopsis, the two bifunctional ATP-grasp kinase/phosphatase enzymes and Vip1 homologs VIH1 and VIH2 catalyze the phosphorylation of InsP_6_ to 1/3-InsP_7_ and 1/3,5-InsP_8_ (*23–25*). Interestingly, VIHs are not only implicated in the generation of PP-InsPs but have also been proposed to contribute to their removal under varying P_i_ concentrations. For instance, the insect-purified C-terminal phosphatase domain of VIH2 hydrolyzed 1-InsP_7_ and 5-InsP_7_ to InsP_6_. Additionally, based on the biochemical activity of yeast Vip1, it has been proposed that low adenylate charge (i.e., ATP/ADP ratios) stimulates the activity of the phosphatase domain of these proteins (*23*). Moreover, fungal Vip1 and Asp1 have been shown to hydrolyze 1,5-InsP_8_ to 5-InsP_7_ (*26–28*). These observations put forward the idea that VIHs might shift their activities to fine-tune energy metabolism and consequently adapt to adverse conditions (*29*).

In addition to local and systemic PSRs, plants form symbiotic associations with microbes to alleviate P_i_ deficiency. Up to 80 % of the land plant species, including major crops such as maize, rice, and wheat, engage in arbuscular mycorrhiza (AM), a mutualistic relationship with *Glomeromycota* fungi (*30*). AM, which originated approximately 450 million years ago, likely contributed to land colonization (*31*, *32*) and influenced climate transition by reducing atmospheric CO_2_ levels, thereby lowering global temperatures and increasing atmospheric oxygen (*33*). AM is a form of endomycorrhiza, where the fungus resides intracellularly within plant roots, establishing a connection between the plant’s root system and the extensive extraradical mycelium of the AM fungus (*31*). Upon symbiotic stimulation, a pre-penetration apparatus forms, guiding the fungus through the root epidermis into deeper cell layers (*34*). Once the cortex is reached, fungal hyphae enter the apoplastic space, where they grow longitudinally and branch out to penetrate the inner cortex cells, forming tree-shaped structures known as arbuscules (*35*, *36*). These arbuscule-containing cells serve as the primary sites for carbon-to-nutrient exchange between the plant and the fungus (*37*). Recent evidence indicates that PSRs and AM symbiosis are interconnected through the SPX/PHR module (*10*, *11*, *38–41*). PHR regulates a network of genes controlling AM development and symbiotic P_i_ uptake (*11*, *39–43*) and specific SPX proteins fine-tune fungal colonization (*10*, *39*, *40*). Moreover, in *Medicago truncatula*, SPX and PHR are involved in arbuscules maintenance (*10*, *41*).

Plants supply their fungal partner with up to 20 % of their photosynthetically fixed carbon in the form of sugars and lipids (*44–49*). In return, AM enhances plant acquisition of P_i_, nitrogen, sulfur, trace elements and water through the specialized hyphal network of the fungus (*50*), while also increasing plant tolerance to various biotic and abiotic stresses (*51*). However, this potential is constrained because even moderate P_i_ levels can systematically inhibit AM colonization, likely as a strategy to conserve carbohydrates (*52*, *53*) and the widespread use of synthetic fertilizers in modern agriculture provides sufficient nutrients to plants, reducing their reliance on AM fungi. A deeper understanding of the molecular mechanisms governing these complex mutualistic associations with phosphate-acquiring microbes, and their integration with nutrient homeostasis, could be instrumental in breeding crop cultivars with improved nutrient use efficiency. Such advancements would enable the cultivation of plants that effectively establish AM symbiosis, reaping all associated short- and long-term benefits while reducing the reliance on mineral fertilizers and minimizing their environmental impact. Here, we demonstrate that *Lotus japonicus* VIH2 is a functional Vip1-type PP-InsP synthase that integrates nutrient homeostasis and symbiosis signaling and regulates AM.

## Results

### *Lotus japonicus* VIH2 is a functional Vip1-type PP-InsP synthase

To study the potential role of PP-InsPs in the regulation of AM and to interrogate the interconnection of P_i_ homeostasis and AM by InsP signaling in *L. japonicus*, we identified the putative *L. japonicus* homolog of *Arabidopsis* VIH2. To assess the enzymatic activity of *L. japonicus* VIH2, we heterologously expressed a long and a short splice variant of its isolated kinase domain as translational fusions with a C-terminal V5-tag in a *Saccharomyces cerevisiae vip1*Δ knock-out mutant, which is unable to grow on 6-azauracil (*54*). While both variants were expressed in yeast (Fig. S1), only the short variant of the isolated *L. japonicus VIH2* kinase domain restored growth of the *S. cerevisiae* mutant on the selection medium (Fig. 1A). In yeast, Vip1 generates 1-InsP_7_ from InsP_6_ and 1,5-InsP_8_ from 5-InsP_7_ (*28*, *55*). While the *S. cerevisiae vip1*Δ knock-out strain no longer produces detectable amounts of InsP_8_ ((*56*); Fig. 1B), a signal corresponding to InsP_8_ was clearly detectable in the normalized strong anion exchange high performance liquid chromatography (SAX-HPLC) profile of InsPs of extracts from [^3^H]-*myo*-inositol-labeled *S. cerevisiae vip1*Δ transformants expressing the short splice variant of the isolated *L. japonicus* VIH2 kinase domain (Fig. 1B). Finally, the short splice variant of the isolated *L. japonicus* VIH2 kinase domain recombinantly expressed and purified from *Escherichia coli* converted 5-InsP_7_ to InsP_8_ *in vitro* (Fig. 1C+D). Our data show that VIH2 is a functional Vip1-type PP-InsP synthase and catalyzes the conversion of InsP_7_ to InsP_8_ in yeast and *in vitro* (Fig. 1).

**Fig. 1.**
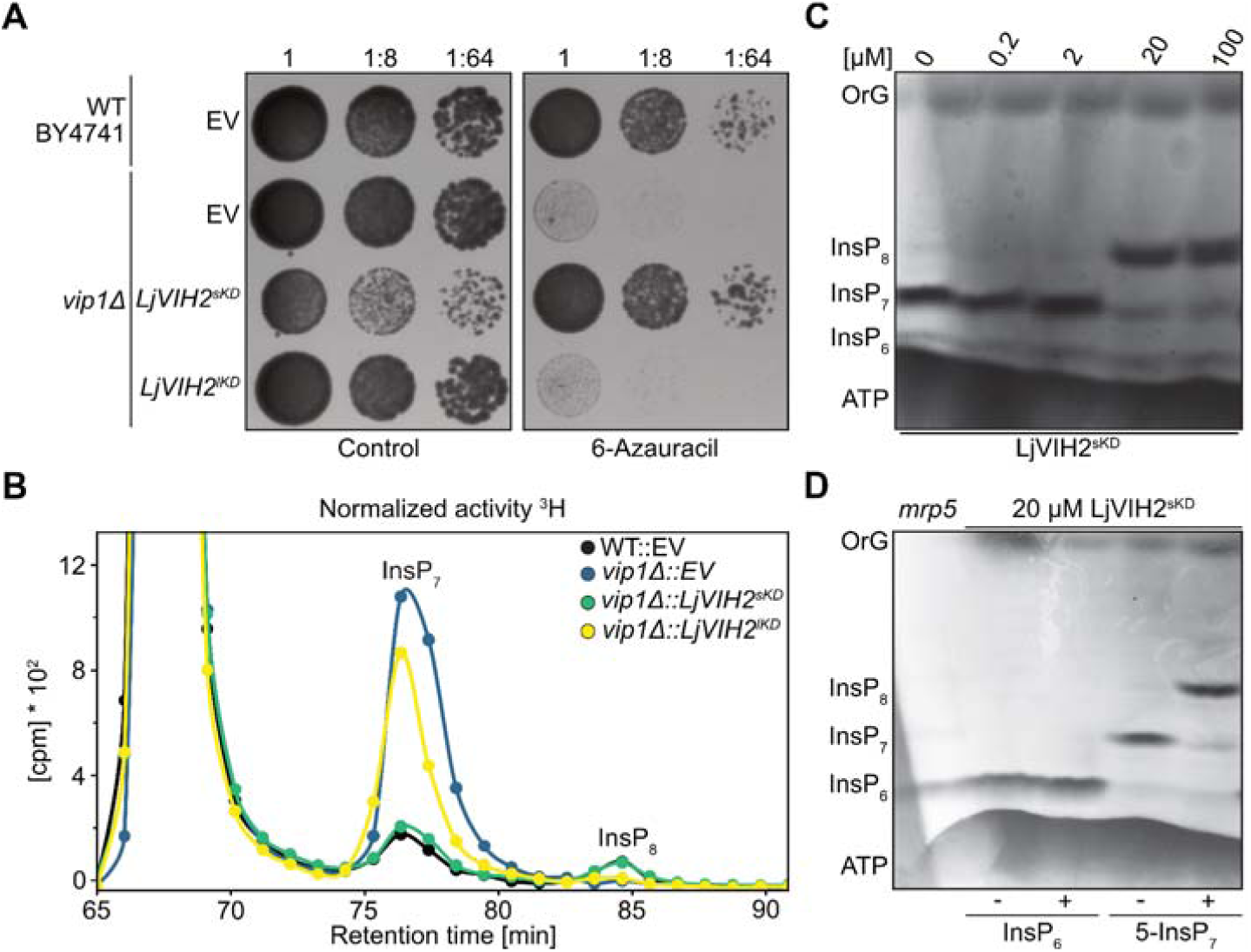
*Lotus japonicus* VIH2 is a functional Vip1-type PP-InsP synthase. The *S. cerevisiae* BY4741 wildtyp (WT) or a *vip1*Δ mutant strain were transformed with the episomal pAG426GPD-*ccdB* empty vector (EV) or with plasmids carrying sequences encoding *S. cerevisiae* Vip1 or either a short (LjVIH2^sKD^) or a long (LjVIH2^lKD^) version of the isolated *L. japonicus* VIH2 kinase domain. **(A)** The indicated yeast strains were spotted in 8-fol serial dilutions onto selective growth media with or without 6-azauracil. Rescue of the *vip1*Δ-associated growth defects on selection medium by *ScVIP1* and *LjVIH2^sKD^* reports Vip1-like activity. **(B)** Normalized HPLC profiles of InsPs of extracts from [^3^H]-*myo*-inositol-labeled yeast transformants either carrying the empty vector (EV) or ectopically expressing a short (*LjVIH2^sKD^*) or a long (*LjVIH2^lKD^*) version of the isolated *L. japonicus* VIH2 kinase domain. Extracts were resolved by SAX-HPLC and fractions were collected each minute for subsequent determination of radioactivity. Expression of *LjVIH2^sKD^* but not of *LjVIH2^lKD^* restored the generation of InsP_8_ in the *vip1*Δ mutant strain. **(C+D)** His_6_-MBP-LjVIH2^sKD^ was expressed in *E. coli* and purified via Ni-NTA resin. **(C)** 0.3 mM 5-InsP_7_ was incubated with increasing concentrations (0–100 µM) of His_6_-MBP-LjVIH2^sKD^ for one hour. **(D)** 0.33 mM InsP_6_ or 5-InsP_7_ were incubated with (+) or without (-) 20 µM recombinant His_6_-MBP-LjVIH2^sKD^ at 22 °C for one hour. Arabidopsis *mrp5* seed extract serves as control. **(C+D)** InsPs were separated by polyacrylamide gel electrophoresis (PAGE) and visualized by toluidine blue staining. cpm, counts per minute; OrG, OrangeG.

### A hydroponic cultivation system to study PSRs and PP-InsP signaling in *Lotus japonicus*

To assess InsP levels, we established a hydroponic system that allows the cultivation of *L. japonicus* under different nutrient regimes (Fig. S2). Lotus plants were grown either with sufficient P_i_ or subjected to P_i_ starvation for eight days. Additionally, P_i_-starved plants were resupplied with P_i_ for 0.5 to 96 hours (Fig. 2), as P_i_ resupply experiments are effective for investigating defects in PP-InsP synthesis by minimizing compensatory effects that occur under steady-state conditions (*24*). The expression of the P_i_ starvation induced (PSI) marker gene *SPX1* was strongly upregulated upon P_i_ starvation and decreased significantly within 0.5 hours of P_i_ resupply (Fig. 2A) confirming successful P_i_ starvation and resupply. InsP_7_ and InsP_8_ were undetectable by PAGE/toluidine blue under both P_i_ sufficiency and deficiency but increased robustly when P_i_-starved plants were resupplied with P_i_ for 48 to 96 hours (Fig. 2B). Quantification of InsP levels by capillary electrophoresis–electrospray ionization-mass spectrometry (CE-ESI-MS) revealed dynamic changes in shoot and root InsP profiles upon P_i_ resupply (*57–60*) (Fig. 2+3, Fig S3+4, 6+7). In shoots, the relative amounts of InsP_8_, 1- and 5- /4/6-InsP_7_ increased more than 25-fold, i.e., from 0–0.2 % reaching 5–7 % of InsP_6_ levels by 48– 96 hours after P_i_ resupply (Fig. 2C; Fig. S3A). Conversely, levels of InsP_3-2_ decreased markedly during this period (Fig. S3A). In roots, relative amounts of InsP_8_, 1- and 5-/4/6-InsP_7_ also rose significantly, albeit to a lower extent, i.e., from 0–0.2 % to 0.5–1.0 % of InsP_6_ by 72–96 hour (Fig. S4A). Additional root InsPs, including InsP_5_ [4/6-OH], 2,3,4,5-InsP_4_, and InsP_3-1_ showed notable increases, while InsP_3-2_ levels declined consistently in roots over the 96-hour period (Fig. S4A). Notably, InsP_6_ levels increased approximately 3-fold in shoots, but remained largely unchanged in roots (Figs S3B+S4B). These results are in line with previous observations in Arabidopsis (*24*) and highlight the suitability of the hydroponics system for studying InsP dynamics in Lotus.

**Fig. 2.**
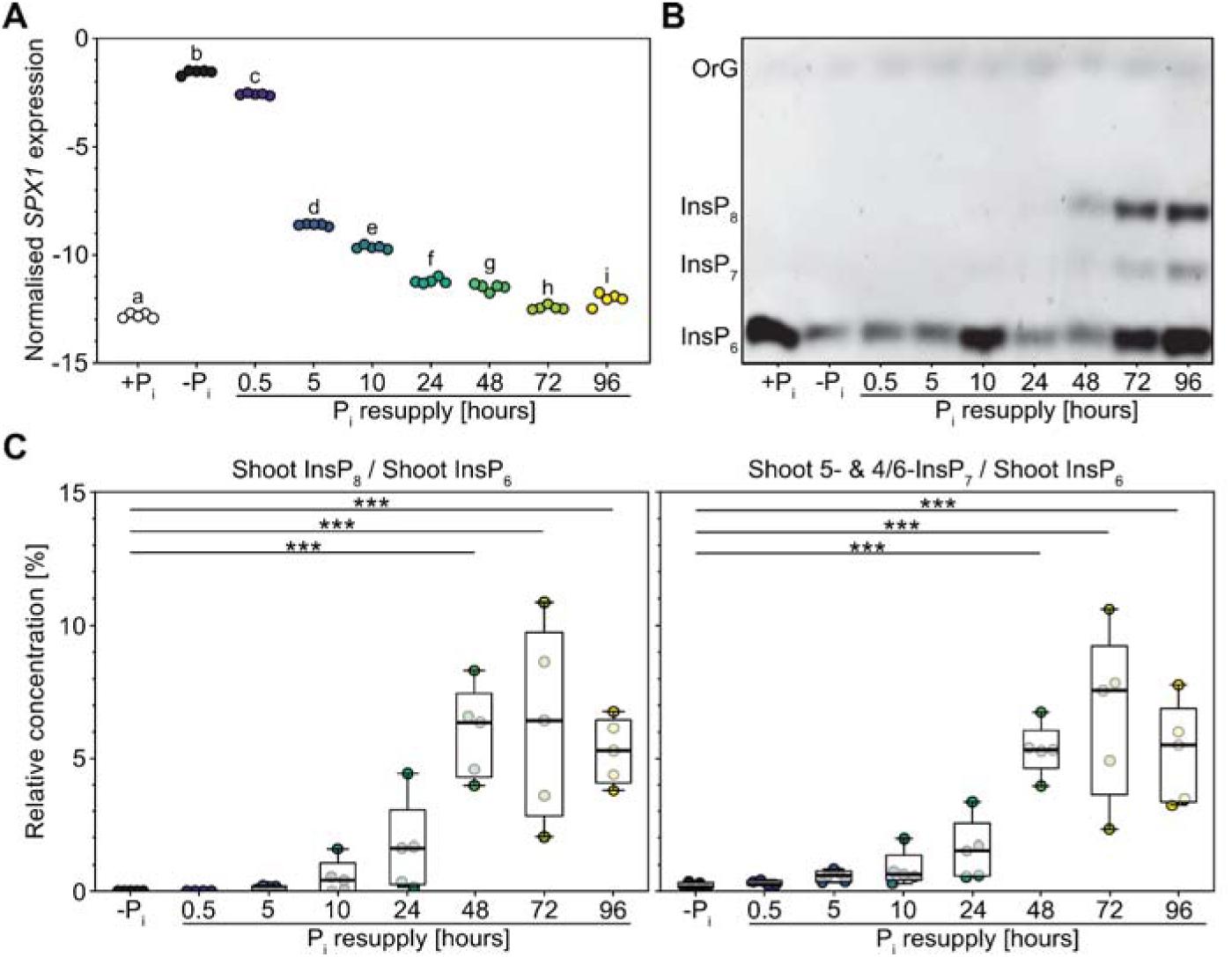
A hydroponic cultivation system to study PSRs and PP-InsP signaling in *Lotus japonicus*. *L. japonicus* wildtype seeds were germinated and seedlings were grown hydroponically in +P_i_ liquid medium containing 150 µM P_i_ for four weeks. Subsequently, plants were either transferred to -P_i_ liquid medium, starved for 8 days (-P_i_), followed by transfer back to +P_i_ liquid medium for P_i_ resupply for 0–96 hours (P_i_ resupply), or kept on +P_i_ liquid medium for the whole time (+P_i_). **(A)** The expression level of the PSI marker gene *SPX1* was analyzed *via* qRT-PCR. n = 5. For statistical analysis, an ordinary one-way ANOVA with Tukey’s multiple comparisons test was performed. Different letters indicate significant differences. **(B+C)** InsPs were enriched by TiO_2_ pull-down and **(B)** resolved by PAGE and visualized by toluidine blue staining or **(C)** quantified *via* CE-ESI-MS analysis. Shoot PP-InsP levels are presented relative to shoot InsP_6_. n = 4–5. For statistical analysis, an ordinary one-way ANOVA with Dunnett’s multiple comparisons test was performed. ***, p ≤ 0.001. OrG, OrangeG.

### *Lotus japonicus vih2* mutants have altered (PP-)InsP levels and show enhanced, constitutive PSRs

To study the potential role of PP-InsPs in the regulation of AM and to interrogate the interconnection of P_i_ homeostasis and AM by InsP signaling in *L. japonicus*, we employed two independent homozygous LORE1 transposon insertion lines (*61*, *62*) in *VIH2* and investigated potential changes in (PP-)InsP levels (Fig. 3A, D-I; Figs S5–7) and associated deregulation of PSRs (Fig. 3B+C). In Arabidopsis, InsP_8_ serves as a proxy for P_i_, and defects in its synthesis result in PSR-related phenotypes (*20*, *23*, *24*). While a clear signal corresponding to InsP_8_ wa previously detected in the normalized SAX-HPLC profile of InsPs of extracts from [^3^H]-*myo*-inositol-labeled *L. japonicus* wildtype seedlings (*63*), we demonstrate that the *L. japonicus vih2-1* and *vih2-2* mutants showed strongly reduced levels of InsP_8_ (Fig. 3A).

**Fig. 3.**
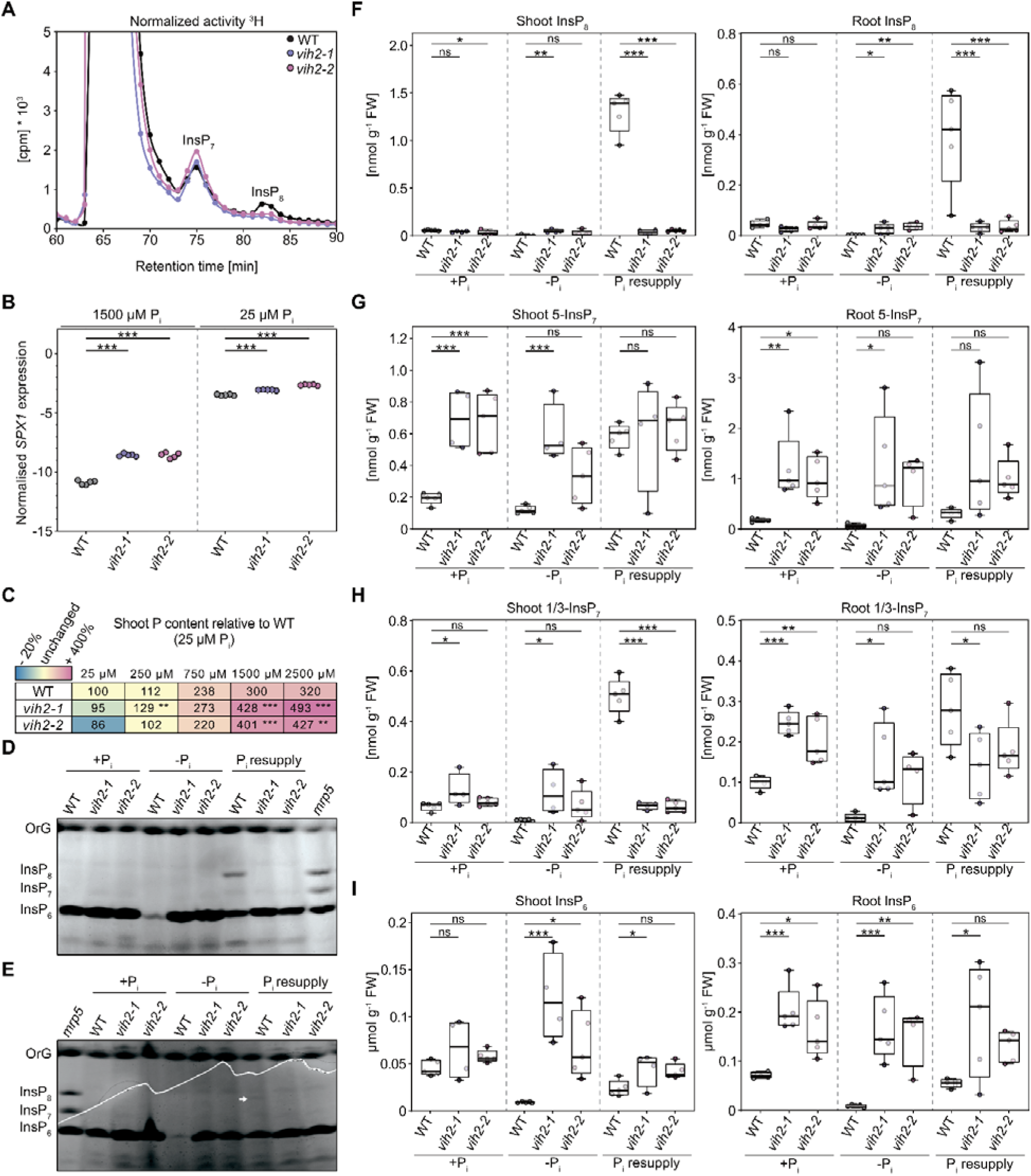
*Lotus japonicus vih2* mutants have altered (PP)-InsP levels and show enhanced, constitutive PSRs. **(A)** *L. japonicus* wildtype (WT) and *vih2* mutant seeds were germinated and subsequently grown for 14 days on water agar plates before transfer to liquid ½ MS media supplemented with 1 % sucrose and 45 µCi of [^3^H]-*myo*-inositol for 5 days. InsPs were extracted and separated by SAX-HPLC. cpm, counts per minute. **(B+C)** Ten-day-old *L. japonicus* wildtype (WT) and *vih2* mutant seedlings were planted in open pots containing 300 mL washed sand (5 seedlings per pot). Plants were fertilized once a week with liquid Lotus cultivation medium containing 25–2500 µM P_i_ and harvested five weeks after planting. **(B)** The expression level of the PSI marker gene *SPX1* was analyzed *vi* qRT-PCR. n = 5. For statistical analysis, an ordinary one-way ANOVA with Dunnett’s multiple comparisons test was performed. ***, p ≤ 0.001. **(C)** Shoot P levels were determined *via* ICP-OES analysis. Data is presented relative to P levels in wildtype shoots grown in the presence of 25 µM P_i_, which were set to 100 %. n = 4–10. Numbers indicate percentages. Stars indicate significant differences of the respective *vih2* mutant line in comparison to the corresponding WT. For statistical analysis, an ordinary one-way ANOVA with Dunnett’s multiple comparisons test was performed. **, p ≤ 0.01; ***, p ≤ 0.001 **(D-I)** *L. japonicus* wildtype (WT) and *vih2* mutant seeds were germinated and seedlings were grown in +P_i_ liquid medium containing 1500 µM P_i_ for four weeks. Subsequently, plants were either transferred to -P_i_ liquid medium, starved for 10 days (-P_i_), followed by transfer back to +P_i_ liquid medium for P_i_ resupply for 72 hours (P_i_ resupply), or kept on +P_i_ liquid medium for the whole time (+P_i_). Shoot and root InsPs were enriched by TiO_2_ pull-down and **(D+E)** resolved by PAGE and visualized by toluidine blue staining, or **(F-I)** quantified *via* CE-ESI-MS analysis. Arrow points to the band corresponding to InsP_8_ in wildtype shoots under P_i_ resupply. n = 4–5. For statistical analysis, an ordinary one-way ANOVA with Dunnett’s multiple comparisons test was performed. *, p ≤ 0.05; **, p ≤ 0.01; ***, p ≤ 0.001. OrG, OrangeG.

As indicated by the expression of the PSI marker gene *SPX1*, PSR in *vih2* mutant lines was indeed constitutively activated at both high (1500 µM), and low (25 µM) external P_i_ (Fig. 3B). However, *vih2-1* and *vih2-2* mutants only accumulated significantly more shoot P than WT plants when grown in the presence of high (1500 or 2500 µM) external P_i_, but not when cultivated with low to medium (25–750 µM) external P_i_ (Fig. 3C). Quantification of InsP levels in hydroponically grown plants by PAGE confirmed a pronounced reduction in InsP_8_ in the shoots and roots of both independent *vih2* mutants under P_i_ resupply conditions (Fig. 3D+E, Fig. S5). This was corroborated by CE-ESI-MS analyses, which additionally revealed a substantial concomitant decrease in shoot 1/3-InsP_7_ under these conditions, along with an increase of InsP_4_, InsP_5_, InsP_6_, 4/6-InsP_7_, and 5-InsP_7_ in shoots under P_i_ deficiency and in roots under P_i_ sufficiency as well as P_i_ deficiency (Fig. 3E-I; Figs S6+S7). Interestingly, while synthesis of InsP_8_ upon P_i_ starvation and resupply in *L. japonicus* shoots is comparable to Arabidopsis shoots, production of InsP_8_ in Lotus roots was markedly increased in comparison to Arabidopsis roots (*24*) (Fig. 3F, Fig. S5).

### Attenuation of VIH2 activity stimulates AM colonization and symbiotic P_i_ uptake at a large range of external P conditions

To investigate whether altered (PP)-InsP synthesis (Fig. 3A, D-F) influences AM colonization, we grew *L. japonicus* wildtype and *vih2* mutant seedlings in a sand-based open pot system and inoculated them with *Rhizophagus irregularis* spores. Thirty days after inoculation, we harvested the root systems and quantified AM colonization. Intriguingly, *L. japonicus vih2* mutants showed significantly increased total, arbuscule and vesicle root length colonization across a large range of external P_i_ concentrations (25–1500 µM Pi), indicating that VIH2-dependent InsPs/PP-InsPs attenuate AM colonization at both high and low external P_i_ levels. (Fig. 4A+B; Fig. S8A-C). This was in line with the expression levels of the two AM marker genes *PT4* and *SbtM1*, which were significantly increased in the *vih2* mutants compared to the wildtype at external P_i_ concentrations of 25–750 µM (Fig. 4C; Fig. S8D). Arbuscules in *vih2* roots were morphologically comparable to those formed in wildtype roots (Fig. 4B, Fig. S8C). Moreover, AM colonization strongly enhanced P acquisition in *vih2* mutant lines: even under low P_i_ conditions (25 µM), both independent *vih2* lines showed a substantial increase in shoot P compared to the wildtype. This effect was further amplified under P_i_-sufficient conditions, resulting in approximately a twofold increase in shoot P compared to wildtype plants, suggesting an additive effect of AM-dependent and AM-independent *vih2*-associated increases in shoot P accumulation (Fig. 4D). Importantly, this AM-dependent effect at low external P_i_ was not restricted to P accumulation alone as we also observed a significant increase of nitrogen, potassium, and magnesium under these conditions (Fig. 4E).

**Fig. 4.**
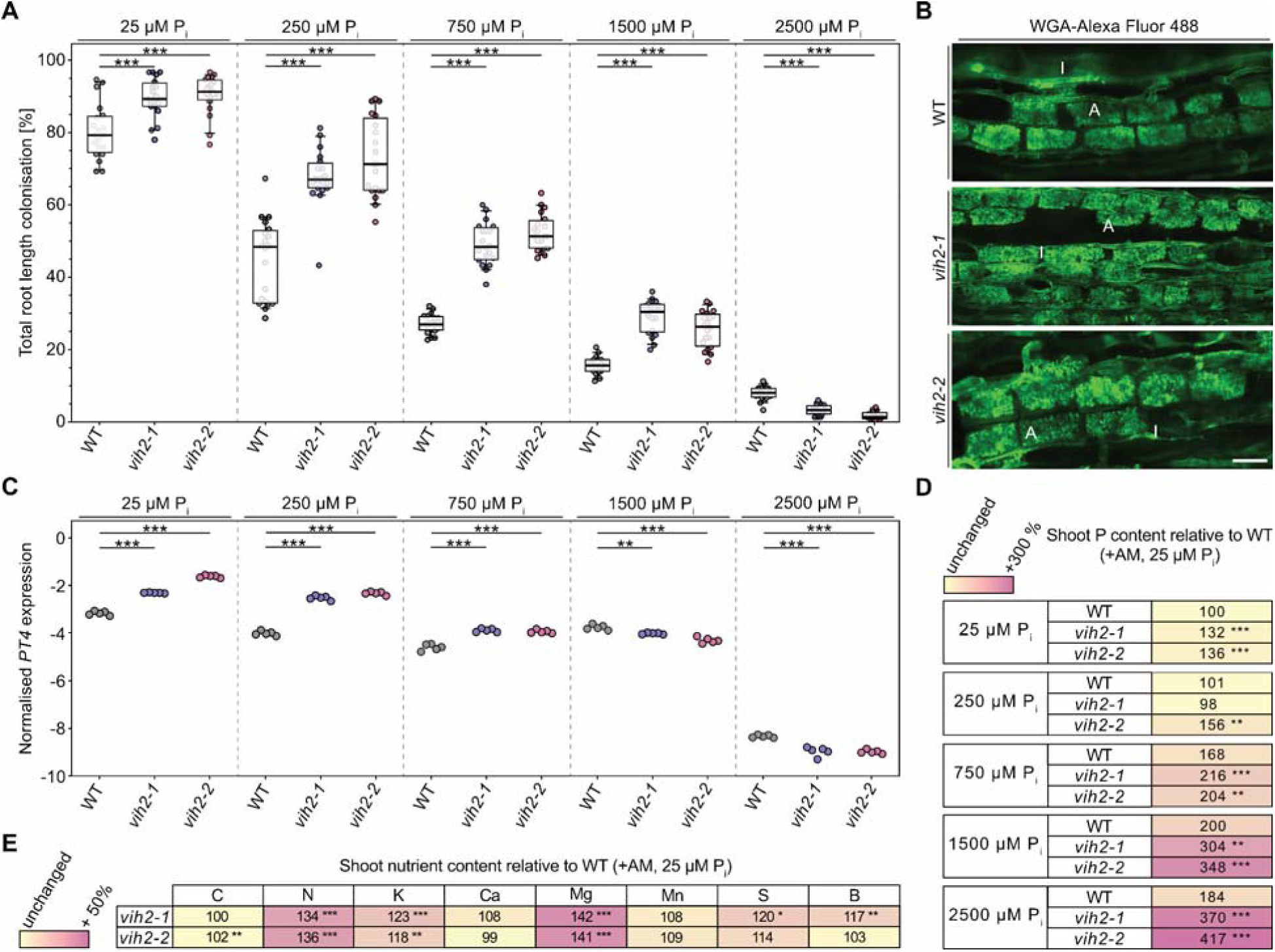
Attenuation of VIH2 activity stimulates AM colonization even under high external P_i_. Ten-day-old *L. japonicus* wildtype (WT) and *vih2* mutant seedlings were planted in open pots containing 300 mL washed sand (5 seedlings per pot) and grown in the presence of *R. irregularis* spore inoculum (Symplanta, Germany; 500 spores per plant). Plants were fertilized once a week with liquid Lotus cultivation medium containing the indicated P_i_ concentrations and harvested 4.5 weeks after planting. **(A)** Roots inoculated with AM fungal spores were staine with ink-vinegar and AM colonization was quantified. n = 20. For statistical analysis, an ordinary one-way ANOVA with Dunnett’s multiple comparisons test was performed. ***, p ≤ 0.001. **(B)** Roots inoculated with AM fungal spores were stained with wheat germ agglutinin conjugated to Alexa Fluor 488 and observed with a confocal microscope. Representative pictures are shown. Bar, 50 µm; WGA, wheat germ agglutinin; A, arbuscule; I, intraradical hypha. **(C)** The expression level of the AM marker gene *PT4* was analyzed in roots inoculated with AM fungal spores *via* qRT-PCR. n = 5. For statistical analysis, an ordinary one-way ANOVA with Dunnett’s multiple comparisons test was performed. **, p ≤ 0.01; ***, p ≤ 0.001. **(D)** Shoot P levels were determined *via* ICP-OES analysis. Data is presented relative to P levels in wildtype shoots of plants inoculated with AM fungal spores and grown in the presence of 25 µM P_i_, which were set to 100 %. n = 5–9. Numbers indicate percentages. Stars indicate significant differences of the respective *vih2* mutant line in comparison to the corresponding WT measurements. For statistical analysis, an ordinary one-way ANOVA with Dunnett’s multiple comparisons test was performed. **, p ≤ 0.01; ***, p ≤ 0.001 **(E)** Shoot nutrient levels were determined *via* ICP-OES or C and N elemental analysis. Data is presented relative to nutrient levels in wildtype shoots of plants inoculated with AM fungal spores grown in the presence of 25 µM P_i_, which were set to 100 %. n = 5–9. Numbers indicate percentages. Stars indicate significant differences of the respective *vih2* mutant line in comparison to the corresponding WT measurements. For statistical analysis, an ordinary one-way ANOVA with Dunnett’s multiple comparisons test was performed. *, p ≤ 0.05; **, p ≤ 0.01; ***, p ≤ 0.001.

To investigate whether the *vih2* mutants were also better colonized by AM fungi than the wildtype at earlier stages of AM development, we investigated total, arbuscule and vesicle root length colonization when the wildtype had a mean total root length colonization of 20 and 40 %, respectively (Fig. S9A+D). Strikingly, already at earlier time points, the *vih2* mutants were significantly better colonized than the wildtype (Fig. S9A+D). The arbuscule morphology wa similar in the roots of all tested genotypes (Fig. S9B+E). Enhanced AM colonization in the *vih2* mutants was accompanied by increased expression levels of the AM marker gene *PT4* (Fig. S9C+F).

### *Lotus japonicus* VIH2 controls AM colonization systemically and locally

We then conducted grafting experiments to determine whether colonization of *L. japonicus* by AM is regulated systemically or locally by VIH2. As expected, a strong increase in AM colonization was observed when roots and shoots of *vih2* plants were self-grafted compared to self-grafted wildtype plants (Fig. 5A, Fig. S10A). Notably, AM colonization also increased to a similar extent when *vih2* shoots were grafted onto wildtype roots, suggesting a strong systemic component in VIH2-dependent AM colonization (Fig. 5A, Fig. S10A). Intriguingly, grafting wildtype shoots onto *vih2* mutant roots also resulted in a robust increase in AM colonization, which was not statistically significantly different from the reciprocal graft, indicating that VIH2 also controls AM colonization locally (Fig. 5A, Fig. S10A). The morphology of arbuscules wa comparable in all tested grafts (Fig. 5B). The expression of the AM marker gene *PT4* largely correlated with total root length colonization (Fig. 5C; Fig. S10B).

**Fig. 5.**
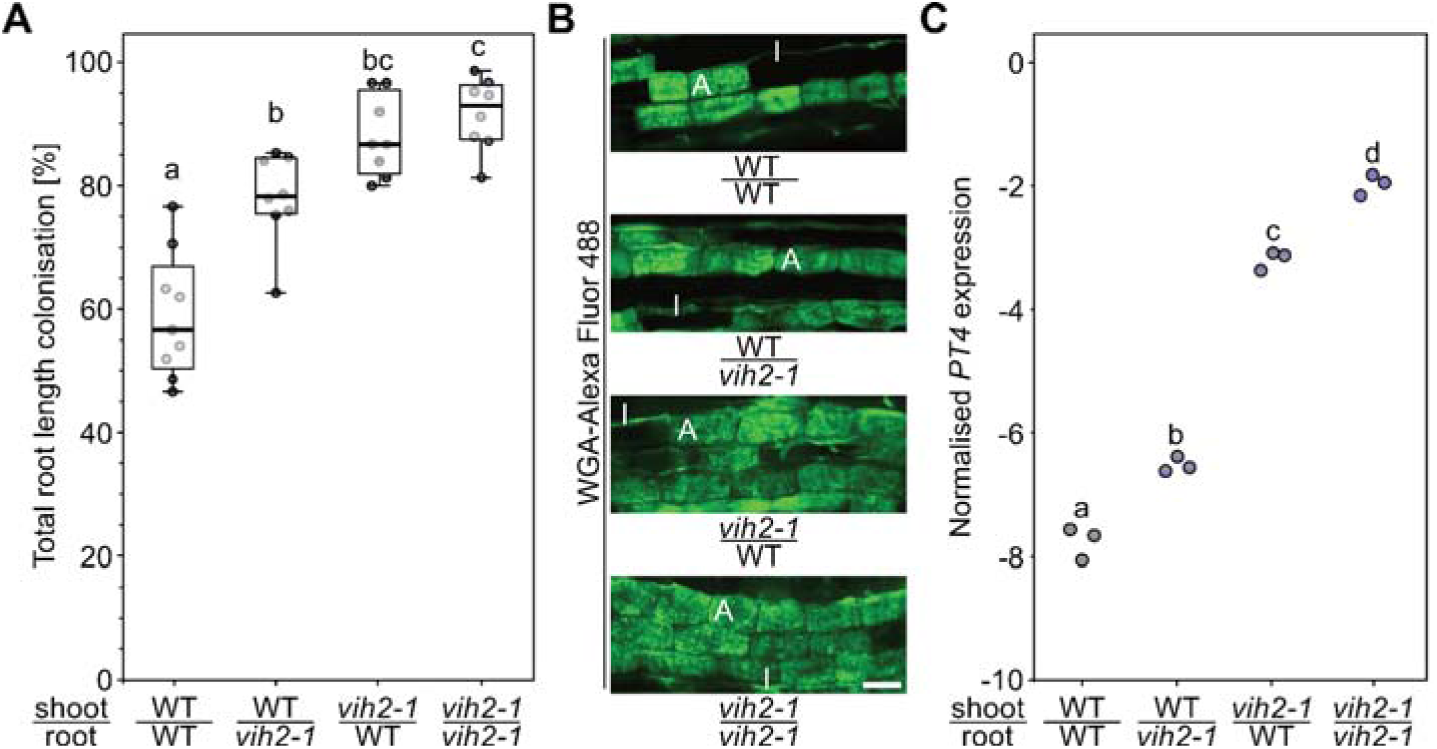
Colonization of *Lotus japonicus* by AM fungi is controlled systemically and locally by *VIH2*. Six-day-old *L. japonicus* wildtype (WT) and *vih2-1* mutant seedlings were self-grafted or reciprocally grafted and grown for three weeks on plates. Subsequently, successful grafts were planted in open pots containing 300 mL washed sand (seedlings per pot) and grown in the presence of *R. irregularis* spore inoculum (Symplanta, Germany; 500 spores per plant). Plants were fertilized once a week with liquid Lotus cultivation medium containing 25 µM P_i_ and harvested five weeks after planting. **(A)** Roots were stained with ink-vinegar and AM colonization was quantified. n = 8–9. For statistical analysis, an ordinary one-way ANOVA with Tukey’s multiple comparisons test was performed. Different letters indicate significant differences. **B)** The expression level of the AM marker gene *PT4* was analyze *via* qRT-PCR. n = 3. For statistical analysis, an ordinary one-way ANOVA with Tukey’s multiple comparisons test was performed. Different letters indicate significant differences. **(C)** Roots were stained with wheat germ agglutinin conjugated to Alexa Fluor 488 and observed with a confocal microscope. Representative pictures are shown. Bar, 5 µm; WGA, wheat germ agglutinin; A, arbuscule; I, intraradical hypha.

## Discussion

Resilience and yield stability are crucial aspects of modern agriculture, particularly as farmers face increasing environmental unpredictability. While maximizing yield has traditionally been the primary goal, there is growing recognition that stable yields over time are equally important. High yield variability poses significant risks to food security and economic stability, making resilience a desirable trait for sustainable farming practices (*64*).

Breeding efforts in the past have primarily focused on easily measurable traits such as disease resistance and drought tolerance. However, incorporating beneficial plant-microbe interactions – like those mediated by AM fungi – into breeding remains challenging as they are influenced by multiple factors, including plant genetics, microbial compatibility, and environmental conditions such as soil properties and nutrient availability (*65*). Plants have evolved mechanisms to suppress AM colonization under high P availability (*52*, *53*), thereby economizing on carbon allocation to the fungal symbiont, a strategy that appears to be ecologically advantageous particularly in natural environments, but not in agriculturally used land. Agricultural practices, including fertilization, soil tillage and monocropping, have been shown to negatively affect AM fungal richness and community composition over the long term (*66*). As a consequence, farmland often exhibits reduced AM fungal diversity and abundance compared to natural ecosystems (*67*, *68*). In consequence, breeding crops capable of sustaining AM colonization under high external P conditions could help preserve AM fungal richness. By ensuring the availability of compatible fungal partners, such crops would support AM colonization across a broader range of phosphate levels and enable the rapid re-establishment of symbiosis under adverse soil conditions. This, in turn, would enhance both resilience and yield stability, addressing the challenges posed by increasingly unpredictable agricultural environments.

Here, we show that Lotus VIH2 functions as a PP-InsP kinase and is largely responsible for the synthesis of InsP_8_, a proxy for P_i_ status in Arabidopsis (*20*, *23*, *24*). We further demonstrate that VIH2 inactivation not only induces PSR (Fig. 3) but also significantly enhances AM colonization, even under high external P_i_ conditions (Fig. 4, Fig. S8, Fig. S9). Notably, in the absence of *R. irregularis*, P_i_ overaccumulation occurred only under high external P_i_ but not at low or medium levels (Fig. 3C), reminiscent of phenotypes in Arabidopsis mutants deficient in InsP_8_ synthesis (*24*). Surprisingly, enhanced AM colonization, as measured by total root colonization, arbuscule abundance, and vesicle formation, was observed consistently across a wide range of external P_i_ concentrations (25–1500 µM) (Fig. 4, Fig. S8, Fig. S9). We speculate that this increased colonization, even at low external P_i_, may stem from a strong local signaling component, independent of systemic P_i_ status, which facilitates AM establishment across varying P_i_ conditions. This phenotype might be driven by elevated VIH2 root activity in Lotus compared to Arabidopsis, as indicated by the stronger InsP_8_ response in Lotus roots following P_i_ resupply to P_i_-starved plants (Fig. 3E+F, Fig. S5), a response absent in Arabidopsis (*24*). Supporting this hypothesis, grafting experiments confirmed that root-specific inactivation of VIH2 is sufficient to enhance AM colonization even at low external P_i_ (Fig. 5A, Fig. S10A).

Our approach complements but also broadens previous strategies to enhance AM colonization by targeting P sensing mechanisms. For example, strong ectopic expression of *PHR* in rice, Lotus (*Ubiquitin* promoter) and Medicago (*35S* promoter) has been shown to promote AM colonization (*11*, *40*, *41*) and improve yield (*11*). However, in Medicago strong ectopic expression of *PHR2* controlled by the *Ubiquitin* promoter resulted in reduced AM development and increased the number of degrading arbuscules (*41*). While effective, such strategies rely on genetically modified organisms (GMOs), which are subject to strict regulatory oversight in many countries. By contrast, our approach could use genome editing to deactivate *VIH* genes, a method that many countries already regulate less stringent or are likely to in the near future (*69*). Also *spx1 spx2 spx3 spx5* quadruple mutants display increased PSR and mycorrhization in rice, while ectopic overexpression of *SPX1* or *SPX2* decreased arbuscule frequency and overall colonization (*40*). Surprisingly, a *spx1 spx3* double mutant in *M. truncatula* exhibited decreased AM colonization, while ectopic overexpression of *SPX1* or *SPX3* resulted in an increase in mycorrhization but also in enhanced arbuscule degradation. While expression of *PHR2* is induced in arbuscule-containing cells in rice but not in Medicago (*40*, *41*), expression of *SPXs* is induced in arbuscule-containing cells in Medicago and tomato but not in rice (*11*, *39*, *40*), suggesting that the SPX-PHR module might be differently wired in different plant species (*10*).

In contrast to previous studies, our strategy builds on modulating signaling molecules like InsP_8_ or 5-InsP_7_ thus providing a more nuanced and adaptable approach, which enables cells to still adjust in response to ATP availability, P_i_ status, and environmental conditions ((*24*); Fig. 3B). Such fine-tuning is not easily achievable in *spx* knockout or *PHR/SPX* overexpression lines, which induce permanent changes in specific regulatory pathways. The finding that local inactivation of VIH2 in the root is sufficient to enhance AM colonization opens further avenues to reduce potential trade-offs associated with manipulating VIH2-dependent PP-InsPs across the entire plant (*25*, *70*). Specifically, ectopic expression of 5-β-phosphate-specific PP-InsP pyrophosphatases, such as PFA-DSPs (*71*, *72*), in root tissues associated with AM colonization might provide a promising direction to calibrate this interaction. This approach would likely reduce unintended consequences caused by altered PP-InsP levels in other plant tissues.

We envisage a range of potential benefits associated with enhanced AM colonization. Improved P-acquisition over a wide range of external P_i_ conditions could help conserve P rock reserves, a globally limited resource, while reducing open-water pollution caused by P runoff and soil erosion. Furthermore, this approach would reduce the input of cadmium and uranium, which are contaminants found in mineral P fertilizers. These heavy metals accumulate in arable soils over time due to P-fertilization, posing health risks to both consumers and farmers (*73*). Further, enhanced AM colonization will likely help plants to better acquire other macro- and micronutrients as well. In support of this, we observed elevated N, K, and Mg levels in the shoots of mycorrhized *vih2* mutant lines, even when grown in our nutrient-rich sand-based substrate (Fig. 4E). Moreover, AM colonization may mitigate the “P–Zn antagonism” phenomenon, where excessive P fertilization reduces zinc (Zn) concentrations in crops contributing to Zn deficiency in humans, an aspect of “hidden hunger” that affects approximately 2 billion people globally (*74–77*). This issue, first described by Bingham and Garber in 1960 (*78*), was recently linked in maize to decreased AM colonization (*79*). Furthermore, AM fungi can improve plant drought tolerance due to the extension of the effective root length by hyphal networks and by improving soil water conductance (*80*), something crops with enhanced AM-colonization might benefit from. Finally AM substantially contributed to global carbon pools and influenced terrestrial biogeochemistry during the Paleozoic climate transition and continues to play a significant role in these processes in the present day (*33*, *81*). An estimated 4 gigatons of CO_2_ equivalents, accounting for about 11 % of global fossil fuel emissions, are transferred annually from plants to AM mycelium (*81*), underscoring the critical role of AM in global climate regulation and the urgent need to integrate AM into future-oriented agricultural practices. Our study presents a novel approach to enhancing AM colonization through the modulation of inositol pyrophosphates, offering a pathway to improve crop resilience and nutrient uptake in changing agricultural environments.

## Materials and Methods

### Molecular cloning, constructs and primers

A detailed description of the constructs used in this study is provided in table S1. A list of primers can be found in table S2. The open reading frames of both (short and long) versions of the *VIH2* kinase domain were amplified from *Lotus japonicus* seedling cDNA by PCR, fused to Gateway compatible *attB*1 and *att*B2 recombination sites and cloned in pDONR221 (Invitrogen) with BP clonase II (Invitrogen). For the expression of recombinant fusion proteins with an N-terminal His_6_-maltose-binding protein (MBP) epitope tag, the ORFs were recombined into the pDest-566 bacterial expression vector (Addgene plasmid #11517 was a gift from Dominic Esposito; http://n2t.net/addgene:11517; RRID:Addgene_11517) by LR clonase II (Invitrogen). For yeast experiments constructs harboring the *VIH2* kinase domains translationally fused to a C-terminal V5 epitope tag were used. The open reading frames harboring the *V5* sequence and Gateway compatible *attB*1 and *att*B2 recombination sites were cloned with BP clonase II in pDONR221 and then recombined into the destination vector pAG426GPD-*ccdB* (*82*) (Addgene plasmid #14156 was a gift from Susan Lindquist; http://n2t.net/addgene:14156; RRID:Addgene_14156) with LR clonase II.

### Plant material

*Lotus japonicus* ecotype Gifu B-129 wildtype and two LORE1 mutant lines (*61*) 30002777 (*vih2-1*) and 30058924 (*vih2-2*) homozygous for a retrotransposon insertion in *L. japonicus VIH2* were used in this study.

### *Lotus japonicus* seed sterilization, hydration, stratification and germination

*L. japonicus* seeds were scarified, surface-sterilized with sterilization solution (1 % sodium hypochlorite, 0.1 % sodium dodecyl sulfate) for 10 minutes, rinsed five times with autoclaved MilliQ water and soaked in autoclaved MilliQ water with end-over-end mixing either for 16 hours at 4 °C or for four hours at room temperature. Subsequently, seeds were stratified on 0.8 % (w/v) plant agar (P1001; Duchefa Biochemie) in square plates (120 mm x 120 mm x 15 mm) for three days at 4 °C in the dark. Plates were transferred to a phytochamber and seeds were germinated for 3 days at 22 °C in the dark.

For SAX-HPLC analyses, *L. japonicus* seeds were scarified, surface-sterilized with 1.2 % sodium hypochlorite for 3 min followed by 70 % ethanol for 3 min and then 100 % ethanol before transfer to filter paper for drying. Subsequently, seeds were stratified on 0.8 % (w/v) bacteriological agar in square plates (120 mm x 120 mm x 15 mm) for three days at 4 °C in the dark before transfer to short day conditions (8 h light at 22 °C, 16 h dark at 20 °C).

### *Lotus japonicus* inoculation with AM fungal spores

After germination, *L. japonicus* seedlings were cultivated for seven days at 22 °C in long day conditions (16 h light, 8 h dark; 160 µmol m^−2^ s^−1^). Five ten-day-old seedlings were transplanted into one pot (9 cm x 9 cm x 9.5 cm) containing 300 mL of washed and autoclaved sand, and each plant was inoculated with 500 spores of *Rhizophagus irregularis* SYMPLANTA-001 research grade (Symplanta; 00101SP). Plants were fertilized once a week with liquid Lotus cultivation medium (25, 250, 750, 1500 or 2500 µM KH_2_PO_4_, 0, 1000, 1750, 2250, or 2475 µM KCl, 5 mM KNO_3_, 50 µM FeNaEDTA, 1 µM MnSO_4_, 0.5 µM ZnSO_4_, 0.2 µM CuSO_4_, 2 µM H_3_BO_4_, 0.1 µM Na_2_MoO_4_, 0.1 µM CoSO_4_, 250 µM MgSO_4_, 1 mM CaCl_2_, 2 mM MES, pH 6.1) and grown at 22 °C in long day conditions (16 h light, 8 h dark; 160 µmol m^−2^ s^−1^). Shoots and roots were harvested at 20 to 35 days post inoculation.

### Ink-vinegar staining and AM quantification

The protocol for ink-vinegar staining of colonized *L. japonicus* roots was adapted from Vierheilig *et al.*, 1998 (*83*). In brief, roots were harvested, washed carefully to remove sand and placed in 2 ml reaction tubes containing 1.5 mL of 10 % KOH. Roots were cleared for 12 hours at 60 °C. KOH was removed and roots were rinsed once with autoclaved MilliQ water and twice with 10 % acetic acid. Roots were covered with 5 % ink-vinegar staining solution (4001; Pelikan) and incubated at 95 °C for 5 minutes. Roots were then rinsed once with autoclaved MilliQ water and once with 5 % acetic acid. Finally, roots were destained in 5 % acetic acid for 20 minutes at room temperature, 5 % acetic acid was replaced with 50 % glycerol and roots were stored at 4 °C until use.

The protocol for the quantification of AM structures was adapted from McGonigle *et al*., 1990 (*84*). In brief, ink-vinegar-stained roots were cut into 1.5-cm-long pieces and for each root system a random selection of 15 root pieces was mounted on a microscope slide and stored at 4 °C until use. Stained roots were analyzed using a Zeiss Axioplan equipped with an Axiocam MRc5 digital camera with 20x magnification. The presence or absence of AM fungal structures (arbuscules, hyphae, vesicles) was scored in ten intersections per root piece resulting in a total of 150 observations per root system. The percentage of a specific AM fungal structure is calculated by dividing the sum of observations in all intersections by the number of intersections (e.g. 75 arbuscules per 150 intersections equals 50 %).

### WGA staining and microscopic analysis

The protocol for WGA staining of colonized *L. japonicus* roots was adapted from Dreher and Yadav *et al*., 2017 (*85*). In brief, roots were harvested and placed in 2 ml reaction tubes containing 1.5 mL 50 % (v/v) ethanol for four hours at room temperature. Roots were then incubated in 20 % KOH for 10 minutes at 90 °C, rinsed once with autoclaved MilliQ water and incubated in 0.1 M HCl for 2 hours at room temperature. Subsequently, roots were transferred to WGA-AlexaFluor488 staining solution (0.2 µg mL^−1^ WGA-Alexa Fluor™ 488 [W11261; ThermoFisher Scientific] in phosphate-buffered saline; 135 mM NaCl, 3 mM KCl, 1.5 mM KH_2_PO_4_, 8 mM Na_2_HPO_4_, pH 7.4) and incubated for minimum four days at room temperature. Stained roots were visualized using a Zeiss LSM 900 confocal microscope with 20x magnification.

### Lotus japonicus grafting

The protocol for grafting of *L. japonicus* was adapted from Sexauer *et al*., 2023 (*86*). In brief, *L. japonicus* seeds were sterilized, hydrated and stratified as described above. For germination, plates were incubated horizontally upside down in a phytochamber for three days at 24 °C in the dark. After germination, plants were transferred to square plates (120 mm x 120 mm x 15mm) containing Lotus cultivation medium supplemented with 750 µM KH_2_PO_4_ and 1 % Noble-Agar (J10907.A1; ThermoFisher Scientific) and cultivated vertically for three days at 21 °C in long day conditions (16 h light, 8 h dark; 140 µmol m^−2^ s^−1^). Seedlings were then cut in the middle of the hypocotyl and shoots were transplanted on root stocks to generate self-grafts and reciprocal grafts between WT and the *vih2-1* mutant. To arrest the fresh grafts, shoots and roots were inserted in sterile silicone tubings (0.76 mm, VWR). Grafts were cultivated on square plates (120 mm x 120 mm x 15mm) containing Lotus cultivation medium supplemented with 750 µM KH_2_PO_4_ and 1 % Noble-Agar (J10907.A1; ThermoFisher Scientific) for seven days at 24 °C in long day conditions (16 h light, 8 h dark; 160 µmol m^−2^ s^−1^) and then subjected to inoculation with AM fungal spores as described above.

### *Lotus japonicus* hydroponic cultivation

*L. japonicus* seeds were sterilized and hydrated as described above. Seeds were germinated in cut-open 1.5 ml reaction tubes filled with rockwool and inserted into flat black boxes (bikapak; Logiflex lid black, 00202201; Logiflex 660 mL box black, 0202233) filled with 250 mL liquid Lotus cultivation medium containing sufficient P_i_ (+P_i_ liquid medium; 1500 µM KH_2_PO_4_, 5 mM KNO_3_, 50 µM FeNaEDTA, 1 µM MnSO_4_, 0.5 µM ZnSO_4_, 0.2 µM CuSO_4_, 2 µM H_3_BO_4_, 0.1 µM Na_2_MoO_4_, 0.1 µM CoSO_4_, 250 µM MgSO_4_, 1 mM CaCl_2_, 2 mM MES, pH 6.1) for 3 days at 22 °C in the dark. Seedlings were further grown in the flat boxes for 17 days and the nutrient solution was exchanged weekly. Plants were then transferred to black 2.5 L buckets (bikapak; round bucket 2.5 L black, 00201989; lid for round bucket 2.5 L black, 00201899) containing 1.8 L +P_i_ liquid medium and grown for seven days. The nutrient solution was exchanged every second day. An air pump (OSAGA, MK-9502) was connected to the system to ensure continuous aeration. Subsequently, plants were transferred to buckets containing P_i_-free liquid Lotus cultivation medium (-P_i_ liquid medium; 1500 µM µM KCl, 5 mM KNO_3_, 50 µM FeNaEDTA, 1 µM MnSO_4_, 0.5 µM ZnSO_4_, 0.2 µM CuSO_4_, 2 µM H_3_BO_4_, 0.1 µM Na_2_MoO_4_, 0.1 µM CoSO_4_, 250 µM MgSO_4_, 1 mM CaCl_2_, 2 mM MES, pH 6.1) and starved for P_i_ for 10 days. Following P_i_ starvation, plants were transferred back to +P_i_ liquid medium for P_i_ resupply for 0.5–96 hours. Plants were maintained at 22 °C in long day conditions (16 h light, 8 h dark; 160 µmol m^−2^ s^−1^) throughout the experiment. Shoots and roots were harvested from plants grown in +P_i_ liquid medium, from P_i_ starved plants and from plants resupplied with P_i_ for the indicated time.

### RNA extraction and cDNA synthesis

Total RNA was extracted from *L. japonicus* tissue using the innuPREP Plant RNA kit (Innuscreen GmbH) according to the manufacturer’s instructions. Lysis Solution PL was applied for root material. The optional DNase I on-column treatment using the innuPREP DNase I Digest kit (Innuscreen GmbH) was performed according to the manufacturer’s instructions to ensure removal of genomic DNA. RNA was eluted in 35 µL RNase-free water and concentration was measured with a spectrophotometer. Absence of genomic DNA was confirmed by PCR. First strand cDNA was synthesized in 20 µL reactions from 500 ng total RNA with the RevertAid First Strand cDNA Synthesis kit (Thermo Scientific) according to the manufacturer’s instructions using the Oligo (dT)_18_ primer. For subsequent gene expression analysis, cDNA was diluted 1:10 in autoclaved MilliQ water.

### Gene expression analysis

qRT-PCR was performed using the PowerUP SYBR Green Master Mix for qPCR (Applied Biosystems) according to the manufacturer’s instructions for a final volume of 10 µl containing 15 ng cDNA and 400 nM of each primer. The reactions were carried out in a QuantStudio 5 real-time PCR system (Applied Biosystems) following the manufacturer’s instructions for the fast-cycling mode. PCR program: 50 °C for 2 minutes, 95 °C for 2 minutes, 40 x (95 °C for 1 second; 60 °C for 30 seconds), 95 °C for 10 s, melt curve 60 °C–95 °C: increment 0.5 °C per 5 s. Expression was normalized to the reference gene *Ubiquitin*. Five biological replicates per sample were analyzed in technical duplicates or triplicates. A primer list can be found in table S2.

### Yeast strains, transformation, growth and complementation (spotting assay)

The BY4741 WT (MATa *his*3Δ *leu*2Δ *met*15Δ *ura*3Δ) and *vip1*Δ (MATa *his*3Δ *leu*2Δ *met*15Δ *ura*3Δ *vip1*Δ*::KanMX*) were obtained from Euroscarf. The yeast cells were transformed with the Li-acetate method (*87*) and grown on 2x CSM-Ura plates for 3 days at 28 °C. Yeast transformants were grown in the respective liquid medium overnight at 28 °C in a rotating wheel before radioactive labeling with [^3^H]-*myo*-inositol (ARC) or performing growth complementation assays. For the latter, OD_600_ from the overnight cultures were measured and the values adjusted to 1.0. Then an 8-fold serial dilution was prepared in a 96-well plate and subsequently 10 μL of each dilution were spotted on selective solid media (*88*) with or without 2 mM 6-azauracil (Sigma-Aldrich; A1757) dissolved in water. After 3 days incubation at 28 °C, pictures were taken with a ChemiDoc^TM^ MP imager (Bio-Rad) using white backlight.

### Radiolabeling of transformed yeast and of *Lotus japonicus* and SAX-HPLC analysis

Transformed yeast were radiolabeled with [^3^H]-*myo*-inositol (ARC) and extracted as published previously (*25*, *89*). *L. japonicus* seedlings were radiolabeled, extracted and analyzed by SAX-HPLC as described before (*63*).

### Yeast protein extraction and immunodetection

Yeast proteins were extracted (*71*) and LjVIH2^KD^ was visualized via immunodetection with a primary mouse anti-V5 antibody (Invitrogen, 1:2000 dilution) followed by a secondary Alexa Fluor™ Plus 680 goat anti-mouse antibody (Invitrogen, 1:10 000 dilution). A rabbit anti-Gal4 antibody (Santa Cruz, 1:1000 dilution) followed by a secondary Alexa Fluor^TM^ Plus 800 goat anti-rabbit antibody (Invitrogen, 1:10 000 dilution) was used as loading control. The fluorescent signals were detected with a ChemiDoc^TM^ MP imager (Bio-Rad).

### Generation of recombinant proteins and *in vitro* kinase assays

His_6_-MBP-fused recombinant VIH2 protein and free His_8_-MBP protein (*25*) were expressed in *E. coli* BL21-CodonPlus(DE3)-RIL cells (Stratagene). The cells were grown in LB medium overnight at 37 °C and diluted 1:500 in 2YT medium the next morning. The cultures were grown at 37 °C for 3 h until protein expression was induced with 100 µM IPTG. They were then transferred to 12 °C and after 3 days the cells were pelleted and resuspended in 15 mL lysis buffer (20 mM Na_2_HPO_4_, 300 mM NaCl, 2 mM DTT, 0.05 mM EDTA, 1 % (v/v) Triton X-100, pH 7.4). 5 mL glass beads (0.1 mm diameter) were added and the tubes vortexed eight times for 1 min, with 1 min on ice between each vortexing step. The samples were centrifuged at 15600 g at 4 °C for 20 min, the supernatant transferred to a fresh 50 mL tube and centrifuged again for 20 min. The supernatant was transferred to 400 μL Ni-NTA agarose beads (Macherey-Nagel), which had previously been washed once with ddH_2_O and twice with lysis buffer. The samples were incubated overnight in an overhead shaker at 4 °C. Next, the samples were centrifuged at 800 g at 4 °C for 1 min, the supernatant was discarded, and the beads were washed three times with binding buffer (20 mM Na_2_HPO_4_, 500 mM NaCl, 25 mM Imidazole, pH 7.4). For elution of the protein, the Ni-NTA resin was incubated three times with 250 μL elution buffer (20 mM Na_2_HPO_4_, 500 mM NaCl, 250 mM Imidazole, pH 7.4) for 5 min and the eluates were collected after centrifugation at 800 g at 4 °C for 1 min. The proteins were desalted with the use of Amicon 50 kDa cut-off centricons (Merck) following the manufacturer’s protocol. For protein storage 20 % glycerol was added. Protein concentrations were estimated by comparison to serum albumin (BSA) standard amounts on a Coomassie blue stained SDS-PAGE.

Kinase reactions were performed by incubating 0.2-100 μM VIH2^sKD^ with 0.33 mM InsP_7_ in 20 mM HEPES (pH 7.5), 5 mM MgCl_2_, 1 mM DTT, 12.5 mM ATP for 1 or 3 h at 22 °C. After incubation InsPs and PP-InsPs were visualized by polyacrylamide gel electrophoresis (PAGE) (*90*).

### Purification of inositol phosphates from *Lotus japonicus,* polyacrylamide gel electrophoresis of inositol phosphates and CE-ESI-MS analysis

Inositol phosphates were extracted with TiO_2_ or Nb_2_O_5_ as indicated and purified as described previously (*91*). Purified inositol phosphates were analyzed by PAGE and by CE-ESI-MS (*24*, *58*, *60*). Enantiomers are not excluded. For InsP_4_ the existence of undefined isomers cannot be excluded.

### Measurement of shoot nutrient content

For nutrient analysis, plant material was dried at 60 °C and ground to powder in a bead mill using agate and steel beads. Approximately 5 mg of the powdered material was analyzed for carbon (C) and nitrogen (N) content using an elemental analyzer (EuroEA 3000, HEKAtech). For the determination of all other reported nutrient concentrations, acid digestions with 50 mg dried plant material and 2.5 mL of concentrated HNO_3_ were performed in a microwave-accelerated reaction system (Mars6, CEM Corporation) followed by analysis with inductively coupled plasma optical emission spectroscopy (ICP-OES; iCAP PRO X Duo, Thermo Fisher Scientific).

### Statistics and data visualization

Statistical analysis and data visualization were performed in GraphPad Prism (version 10.0.3 for Windows, GraphPad Software, Boston, Massachusetts USA, www.graphpad.com) and Affinity Designer (version 2.5.5 for Windows, https://affinity.serif.com/de/designer/). Boxplots together with individual data points were used to display data in Figs. 2C, 3F–I, 4A, 5A, S3, S4, S6, S7, S8A+B, S9A+D, and S10A (black line, median; box, 25^th^ to 75^th^ percentiles; whiskers, 10^th^ to 90^th^ percentiles. Individual data points only were plotted in Figs. 2A, 3B, 4C, 5C, S8D, S9D+F, and S10B. Statistical tests applied are stated in the figure legends.

## Acknowledgments

We thank Christine Wagner for initial work with the hydroponic cultivation system, Katja Baumann-Kaschig (Department of Molecular Signal Processing, Leibniz Institute of Plant Biochemistry), Brigitte Ueberbach, Li Schlüter, Angelika Glogau, Nur Gömec and Anna M. Frentzen (Department of Plant Nutrition, Institute of Crop Science and Resource Conservation, University of Bonn) for excellent technical assistance, and Prof. Dr. Bettina Hause and Hagen Stellmach (Department of Cell and Metabolic Biology, Leibniz Institute of Plant Biochemistry) for support with microscopy.

## Author contributions

Conceptualization: MKRL, GS, VG

Methodology: MKRL, GS, HJJ, VG, KR

Formal analysis: MKRL, VG

Investigation: KR, VG, ML, MS, PG, CMMG, UAJ, GL, MH

Visualization: MKRL

Funding acquisition: MKRL, GS, HJJ

Project administration: MKRL, GS, VG

Supervision: MKRL, GS, HJJ, VG

Original draft: MKRL, GS

Review & editing: MKRL, GS, VG, PG, KR, ML, UAJ, HJJ

## Data and materials availability

All data and materials that support the findings of this publication are available within this paper or its Supplementary Materials or are available from the corresponding authors upon reasonable request.

## Supplementary Material

**Fig. S1.**
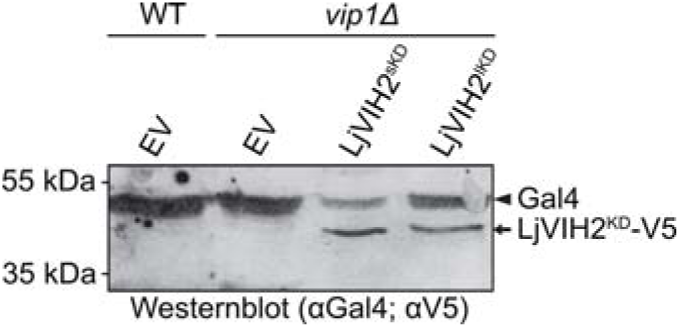
*Lotus japonicus* VIH2 is expressed in the *Saccharomyces cerevisiae vip1*Δknock-out mutant strain. The *S. cerevisiae* wildtype or *vip1*Δ mutant strain was transformed with the episomal pAG426GPD-*ccdB* empt vector (EV) or with plasmids carrying sequences encoding either a short (LjVIH2^sKD^) or a long (LjVIH2^lKD^) versio of the isolated *L. japonicus* VIH2 kinase domain fused to a C-terminal V5-tag. Gal4 served as loading control.

**Fig. S2.**
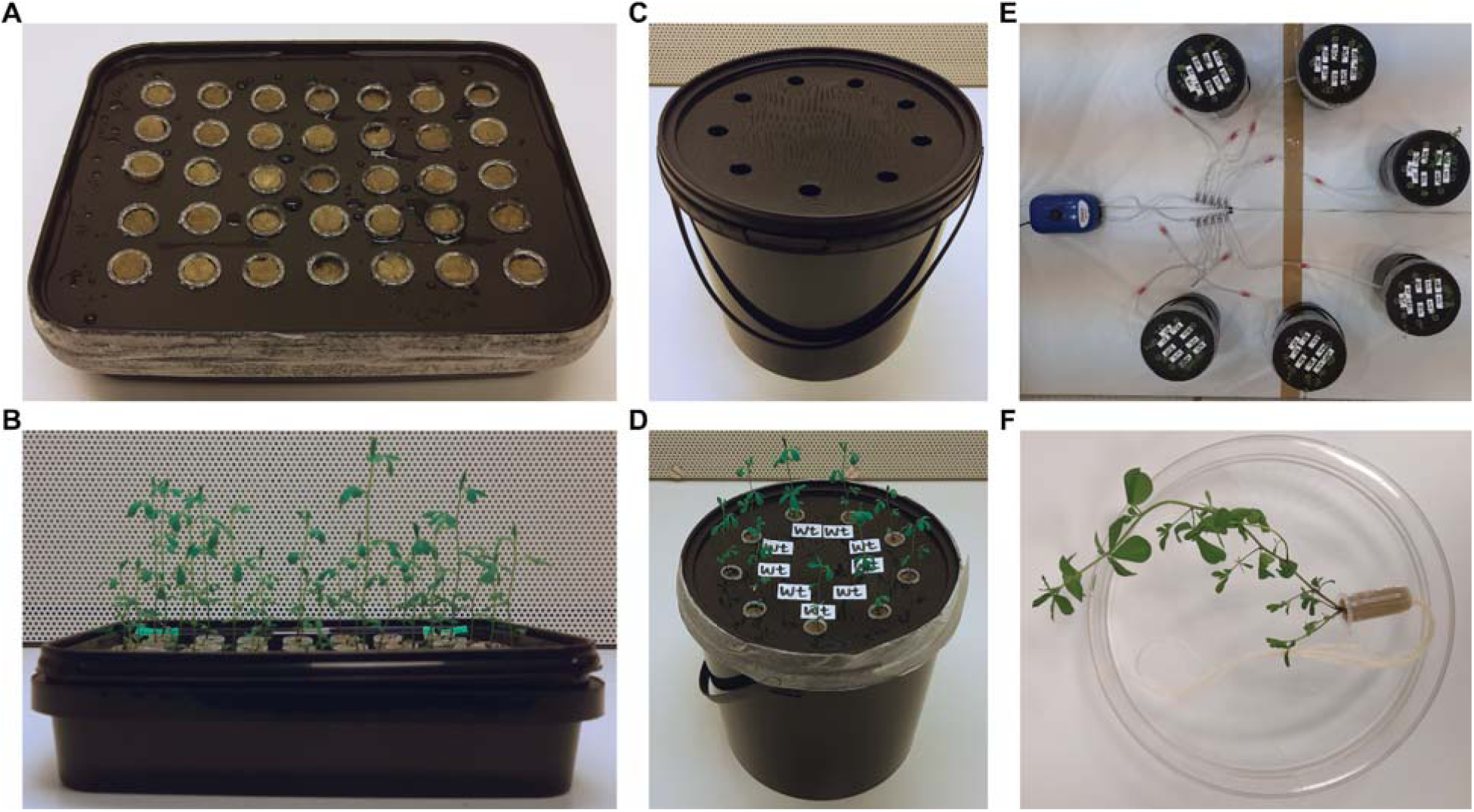
A hydroponic cultivation system to study PSRs and InsP signaling in *Lotus japonicus*. (A+B) *L. japonicus* seeds were germinated in cut-open 1.5 mL reaction tubes filled with rockwool and placed into flat blac boxes (bikapak; Logiflex lid black, 00202201; Logiflex 660 mL box black, 0202233) containing 250 mL liquid Lotus cultivation medium. **(C + D)** Once the root systems reached the desired length, plants were transferred t black 2.5 L buckets (bikapak; round bucket 2.5 L black, 00201989; lid for round bucket 2.5 L black, 00201899) containing 1.8 L liquid Lotus cultivation medium. **(E)** Buckets were connected to an air pump (OSAGA, MK-9502) to ensure continuous aeration. **(F)** Reaction tubes were removed from the bucket lids for harvesting.

**Fig. S3.**
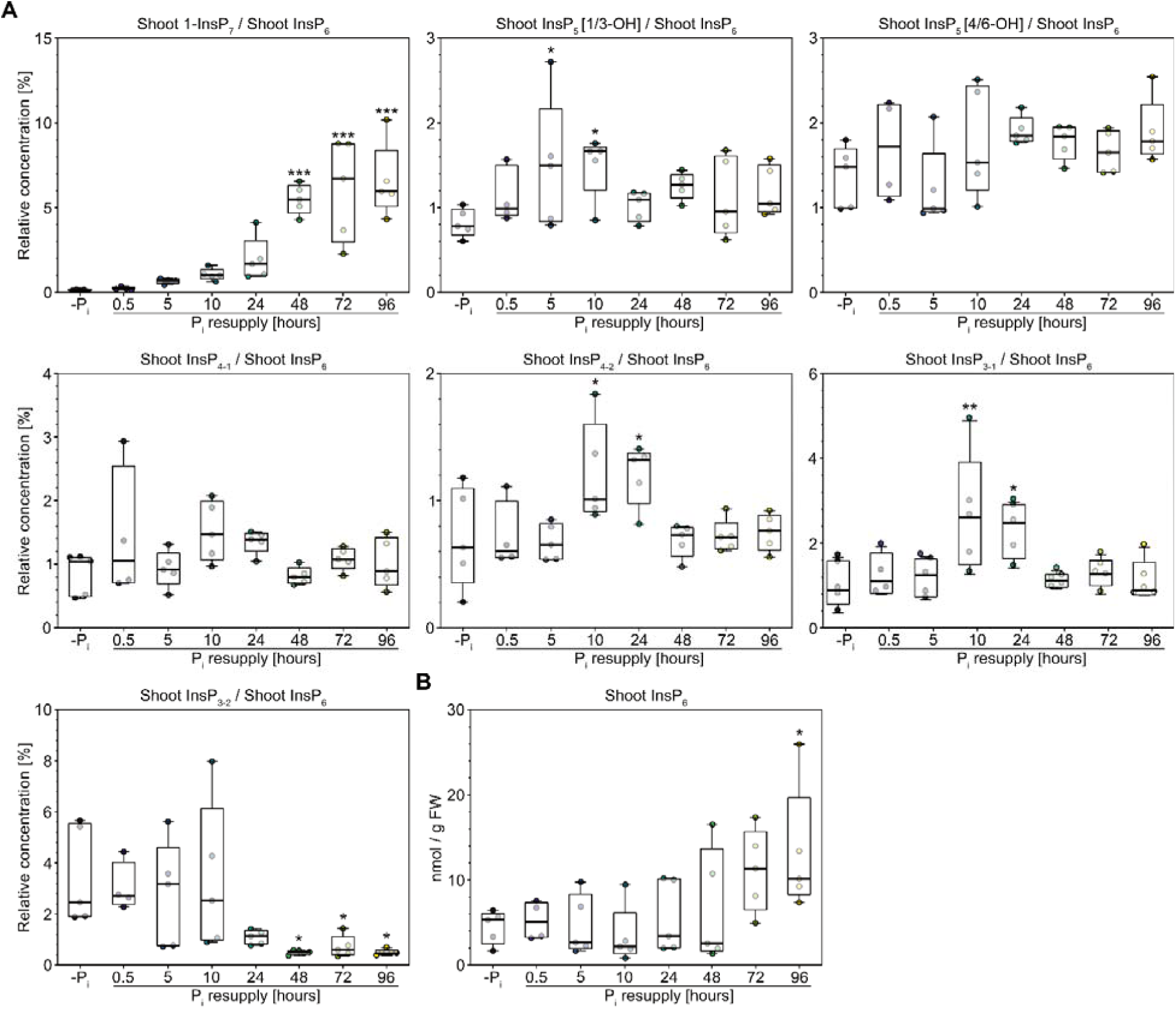
Changes in shoot InsP levels in hydroponically grown *Lotus japonicus* upon phosphate starvation and resupply. *L. japonicus* wildtype seeds were germinated and seedlings were grown in +P_i_ liquid medium containing 1500 µM P_i_ for four weeks. Subsequently, plants were transferred to -P_i_ liquid medium, starved for 8 days (-P_i_), followed by transfer back to +P_i_ liquid medium for P_i_ resupply for 0.5 to 96 hours (Pi resupply). Shoot InsPs were enriched by TiO_2_ pull-down and quantified *via* CE-ESI-MS analysis. **(A)** Shoot InsP levels are presented relative t shoot InsP_6_. InsP_4-1_ contains 1,4,5,6-InsP_4_, whereas InsP_4-2_ contains 2,3,4,5-InsP_4_ but the potential presence of undefined isomers cannot be excluded. InsP_3-1_ represents an inseparable mixture that likely includes 1,2,3-InsP_3_, 3,4,5-InsP_3_, and 1,2,6-InsP_3_, whereas InsP_3-2_ comprises an inseparable mixture potentially containing 1,3,4-InsP_3_, 1,4,5-InsP_3_, and 1,4,6-InsP_3_. **(B)** Shoot InsP_6_ levels. n = 4–5. For statistical analysis, an ordinary one-way ANOVA with Dunnett’s multiple comparisons test was performed. Every time point was compared to -P_i_. *, p ≤ 0.05; **, p ≤ 0.01; ***, p ≤ 0.001.

**Fig. S4.**
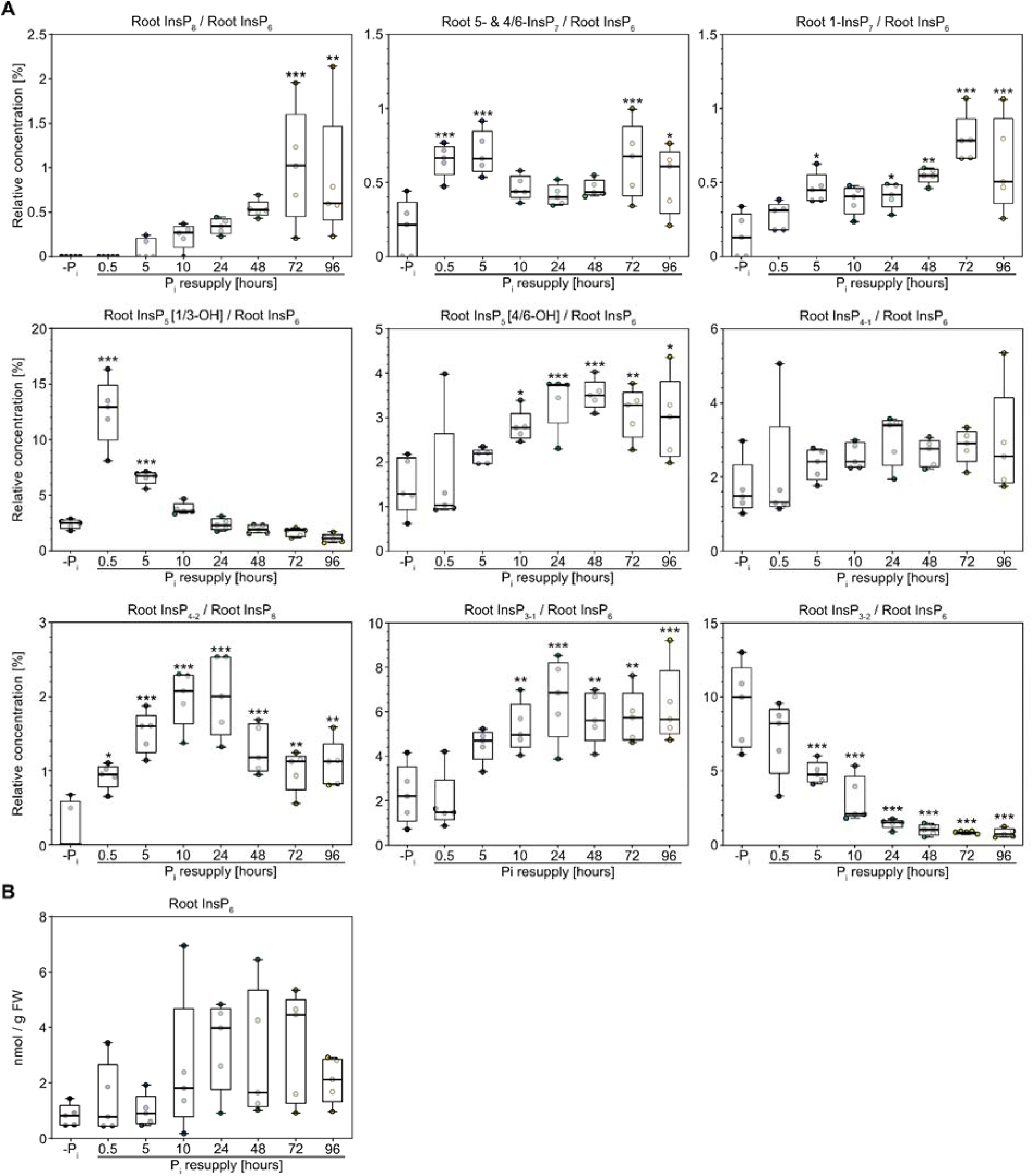
Changes in root InsP levels in hydroponically grown *Lotus japonicus* upon phosphate starvation and resupply. *L. japonicus* wildtype seeds were germinated and seedlings were grown in +P_i_ liquid medium containin 1500 µM P_i_ for four weeks. Subsequently, plants were transferred to -P_i_ liquid medium, starved for 8 days (-P_i_), followed by transfer back to +P_i_ liquid medium for P_i_ resupply for 0.5 to 96 hours (P_i_ resupply). Root InsPs were enriched by TiO_2_ pull-down and quantified *via* CE-ESI-MS analysis. **(A)** Root InsP levels are presented relative t root InsP_6_. InsP_4-1_ contains 1,4,5,6-InsP_4_, whereas InsP_4-2_ contains 2,3,4,5-InsP_4_ but the potential presence of undefined isomers cannot be excluded InsP_3-1_ represents an inseparable mixture that likely includes 1,2,3-InsP_3_, 3,4,5-InsP_3_, and 1,2,6-InsP_3_, whereas InsP_3-2_ comprises an inseparable mixture potentially containing 1,3,4-InsP_3_, 1,4,5-InsP_3_, and 1,4,6-InsP_3_. **(B)** Root InsP_6_ levels. n = 5. For statistical analysis, an ordinary one-way ANOVA wit Dunnett’s multiple comparisons test was performed. Every time point was compared to -P_i_. *, p ≤ 0.05; **, p ≤ 0.01; ***, p ≤ 0.001.

**Fig. S5.**
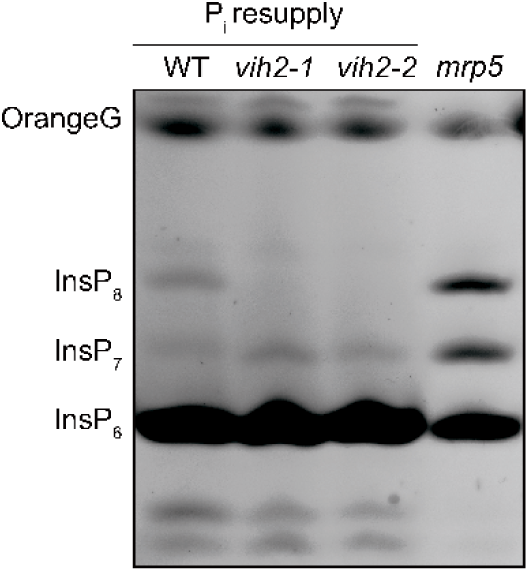
*Lotus japonicus vih2* mutants have altered PP-InsP levels. *L. japonicus* wildtype (WT) and *vih2* mutant seeds were germinated and seedlings were grown in +P_i_ liquid medium containing 1500 µM P_i_ for four weeks. Subsequently, plants were transferred to -P_i_ liquid medium, starved for 10 days, followed by transfer back to +P_i_ liquid medium for P_i_ resupply for 72 hours (P_i_ resupply). Root InsPs were enriched by Nb_2_O_5_ pull-down and **r**esolved by PAGE and visualized by toluidine blue and subsequent DAPI staining. OrG, OrangeG.

**Fig. S6.**
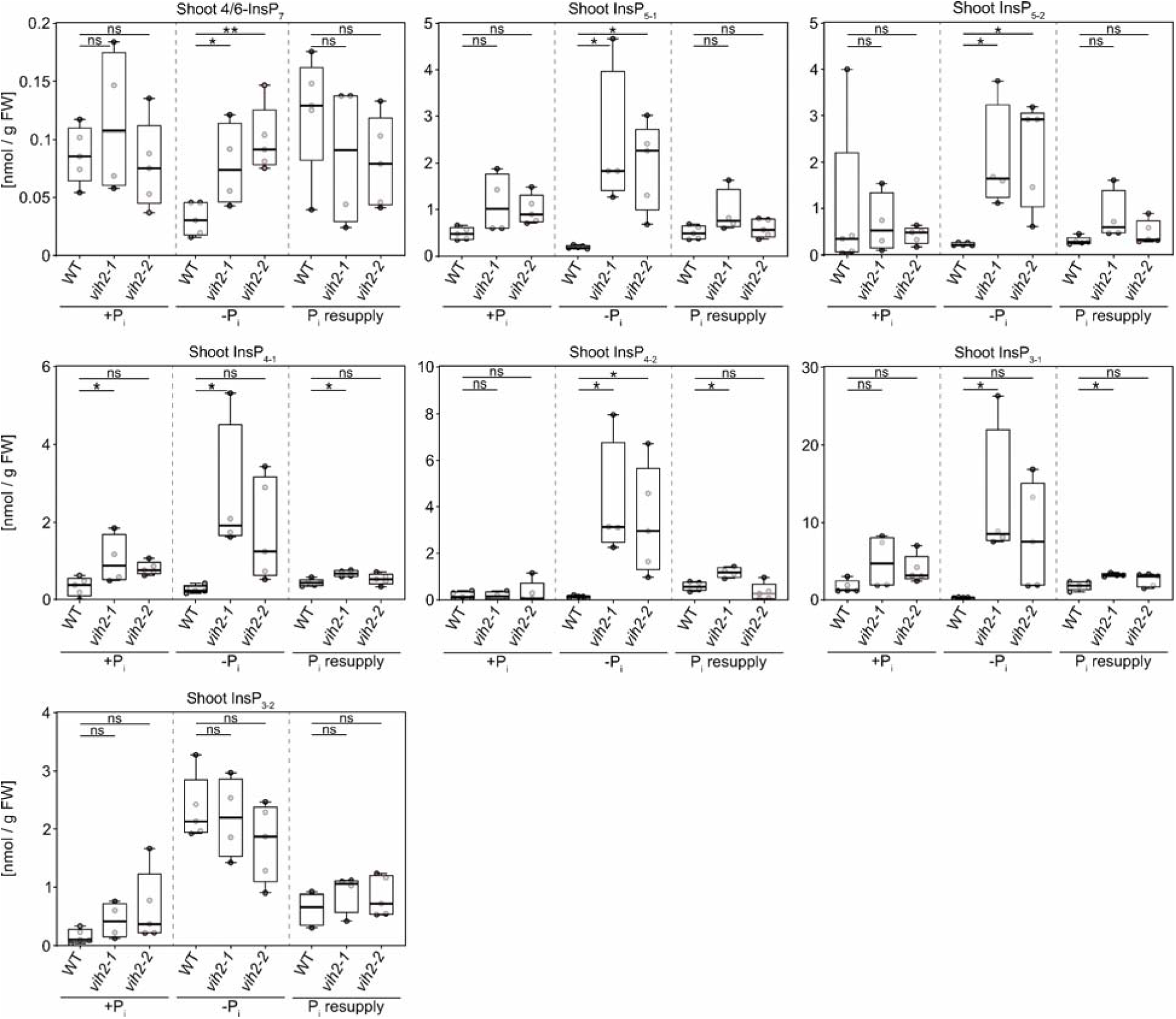
*Lotus japonicus vih2* mutants have altered shoot InsP levels. *L. japonicus* wildtype (WT) and *vih* mutant seeds were germinated and seedlings were grown in +P_i_ liquid medium containing 1500 µM P_i_ for four weeks. Subsequently, plants were either transferred to -P_i_ liquid medium, starved for 10 days (-P_i_), followed by transfer back to +P_i_ liquid medium for P_i_ resupply for 72 hours (P_i_ resupply), or kept on +P_i_ liquid medium for the whole time (+P_i_). Shoot InsPs were enriched by Nb_2_O_5_ pull-down and quantified *via* CE-ESI-MS analysis. InsP_5-1_ may include InsP_5_ [4/6-OH] and InsP_5_ [5-OH], while InsP_5-2_ refers to InsP_5_ [2-OH] and InsP_5_ [1/3-OH]. InsP_4-1_ contains 1,4,5,6-InsP_4_, whereas InsP_4-2_ contains 2,3,4,5-InsP_4_ but the potential presence of undefined isomers cannot be excluded. InsP_3-1_ represents an inseparable mixture that likely includes 1,2,3-InsP_3_, 3,4,5-InsP_3_, and 1,2,6-InsP_3_, whereas InsP_3-2_ comprises an inseparable mixture potentially containing 1,3,4-InsP_3_, 1,4,5-InsP_3_, and 1,4,6-InsP_3_. = 4–5. For statistical analysis, an ordinary one-way ANOVA with Dunnett’s multiple comparisons test was performed. *, p ≤ 0.05; **, p ≤ 0.01; ***, p ≤ 0.001.

**Fig. S7.**
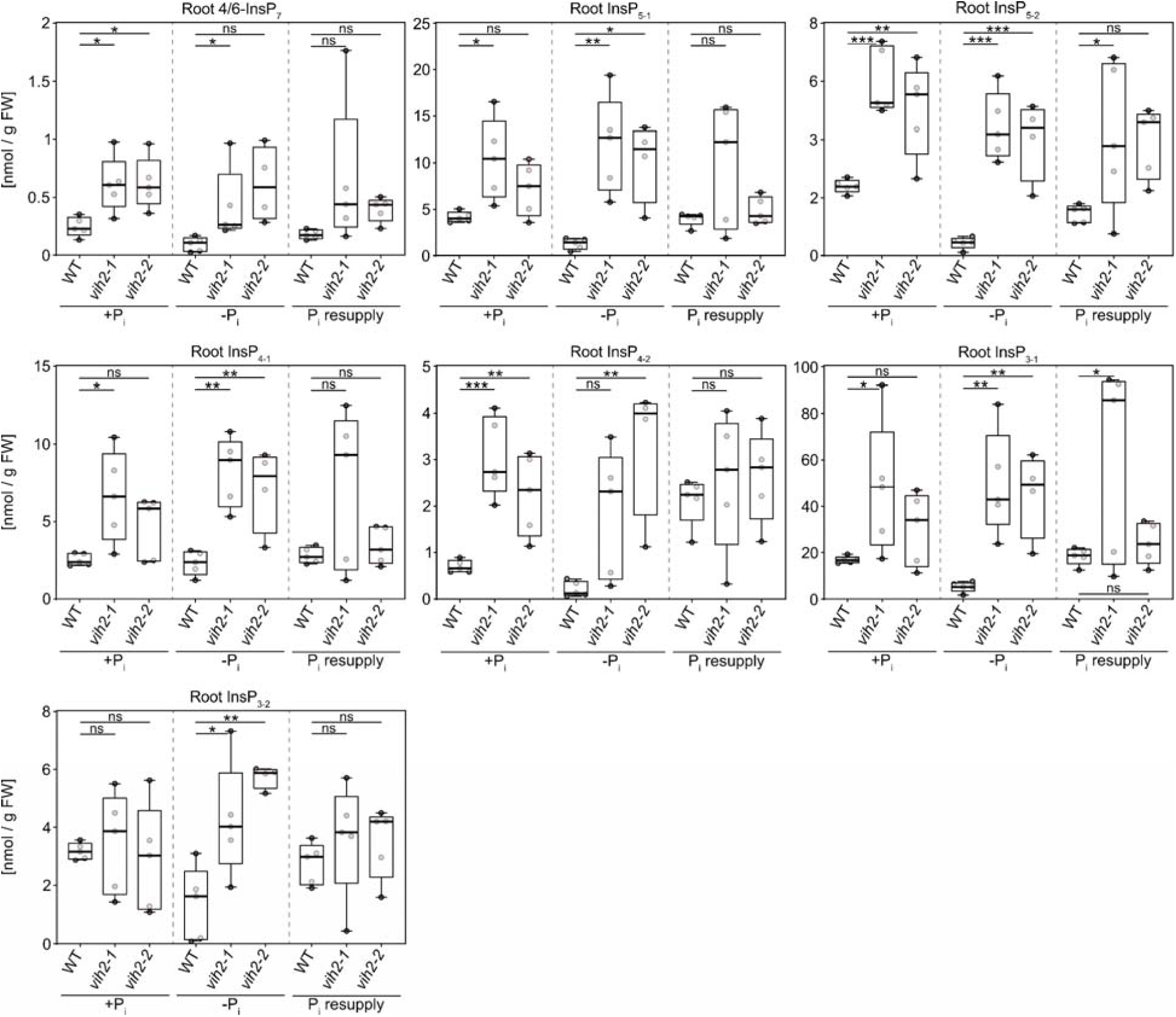
*Lotus japonicus vih2* mutants have altered root InsP levels. *L. japonicus* wildtype (WT) and *vih2* mutant seeds were germinated and seedlings were grown in +P_i_ liquid medium containing 1500 µM P_i_ for four weeks. Subsequently, plants were either transferred to -P_i_ liquid medium, starved for 10 days (-P_i_), followed by transfer back to +P_i_ liquid medium for P_i_ resupply for 72 hours (P_i_ resupply), or kept on +P_i_ liquid medium for the whole time (+P_i_). Root InsPs were enriched by Nb_2_O_5_ pull-down and quantified *via* CE-ESI-MS analysis. InsP_5-1_ ma include InsP_5_ [4/6-OH] and InsP_5_ [5-OH], while InsP_5-2_ refers to InsP_5_ [2-OH] and InsP_5_ [1/3-OH]. InsP_4-1_ contains 1,4,5,6-InsP_4_, whereas InsP_4-2_ contains 2,3,4,5-InsP_4_ but the potential presence of undefined isomers cannot b excluded. InsP_3-1_ represents an inseparable mixture that likely includes 1,2,3-InsP_3_, 3,4,5-InsP_3_, and 1,2,6-InsP_3_, whereas InsP_3-2_ comprises an inseparable mixture potentially containing 1,3,4-InsP_3_, 1,4,5-InsP_3_, and 1,4,6-InsP_3_. = 4–5. For statistical analysis, an ordinary one-way ANOVA with Dunnett’s multiple comparisons test was performed. *, p ≤ 0.05; **, p ≤ 0.01; ***, p ≤ 0.001.

**Fig. S8.**
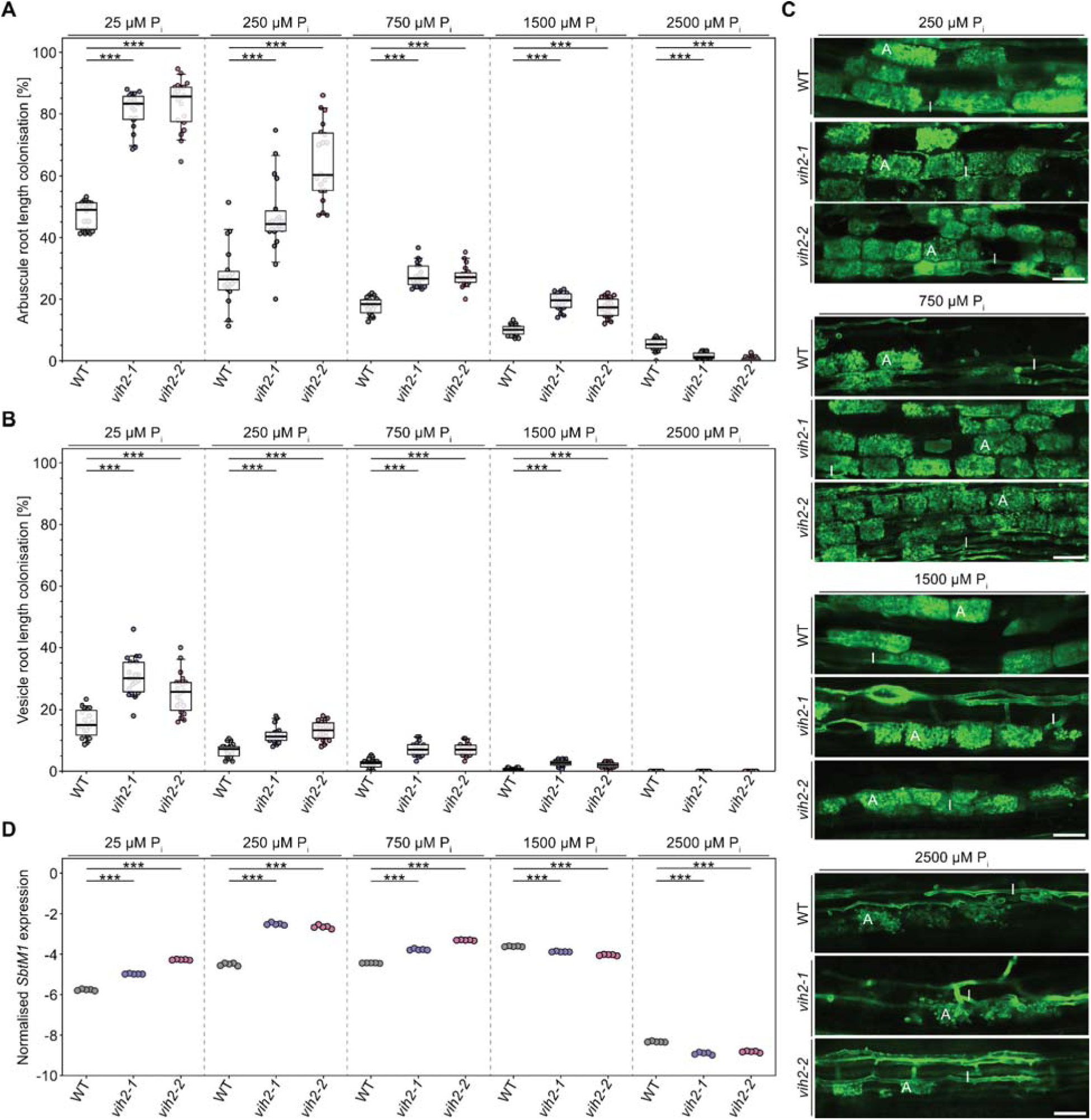
*Lotus japonicus vih2* mutants are significantly better colonized by AM fungi than wildtype plants. Ten-day-old *L. japonicus* wildtype (WT) and *vih2* mutant seedlings were planted in open pots containing 300 mL washed sand (5 seedlings per pot) and inoculated with *R. irregularis* spores (Symplanta, Germany; 500 spores per plant). Plants were fertilized once a week with liquid Lotus cultivation medium containing the indicated P_i_ concentrations and harvested 4.5 weeks after planting. Roots were stained with ink-vinegar and arbuscules **(A)** an vesicles **(B)** were quantified. n = 20. For statistical analysis, an ordinary one-way ANOVA with Dunnett’s multiple comparisons test was performed. ***, p ≤ 0.001. **(C)** Roots were stained with wheat germ agglutinin conjugated to Alexa Fluor 488 and observed with a confocal microscope. Representative pictures are shown. Bar, 50 µm; WGA, wheat germ agglutinin; A, arbuscule; I, intraradical hypha. **(D)** The expression level of the AM marker gene *SbtM* was analyzed *via* qRT-PCR. n = 5. For statistical analysis, an ordinary one-way ANOVA with Dunnett’s multiple comparisons test was performed. ***, p ≤ 0.001.

**Fig. S9.**
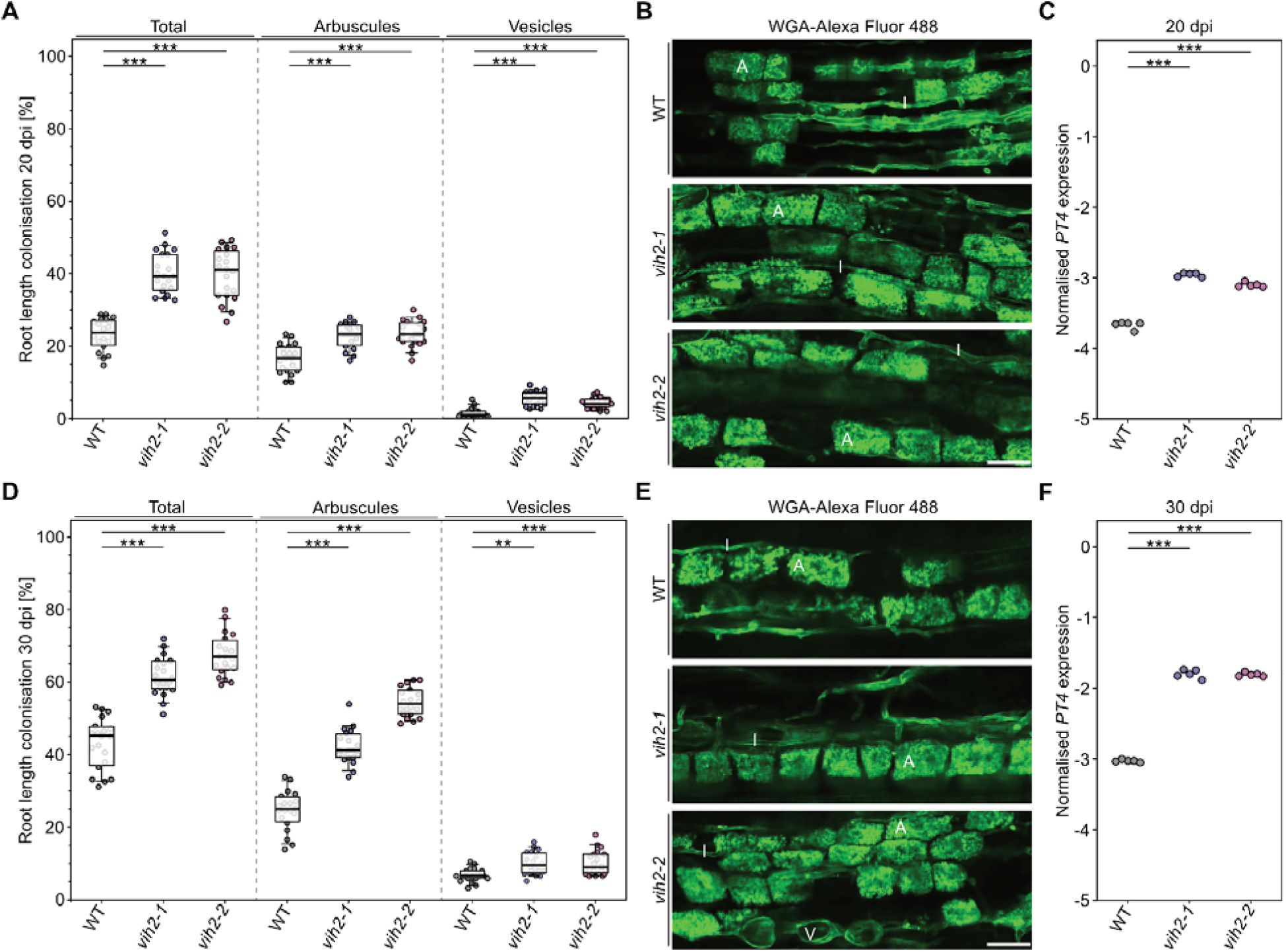
*Lotus japonicus vih2* mutants are significantly better colonized by AM fungi than wildtype plants. Ten-day-old *L. japonicus* wildtype (WT) and *vih2* mutant seedlings were planted in open pots containing 300 mL washed sand (5 seedlings per pot) and grown in the presence of *R. irregularis* spore inoculum (Symplanta, Germany; 500 spores per plant). Plants were fertilized once a week with liquid Lotus cultivation medium containin 25 µM P_i_ harvested 20 **(A-C)** or 30 dpi **(D-F)** after planting. **(A+D)** Roots were stained with ink-vinegar and AM colonization was quantified. n = 20. For statistical analysis, an ordinary one-way ANOVA with Dunnett’s multiple comparisons test was performed. **, p ≤ 0.01; ***, p ≤ 0.001. **(B+E)** Roots were stained with wheat germ agglutinin conjugated to Alexa Fluor 488 and observed with a confocal microscope. Representative pictures are shown. Bar, 5 µm; WGA, wheat germ agglutinin; V, vesicles; A, arbuscule; I, intraradical hypha. **(C+F)** The expression level of the AM marker gene *PT4* was analyzed *via* qRT-PCR. n = 5. For statistical analysis, an ordinary one-way ANOVA with Dunnett’s multiple comparisons test was performed. ***, p ≤ 0.001.

**Fig. S10.**
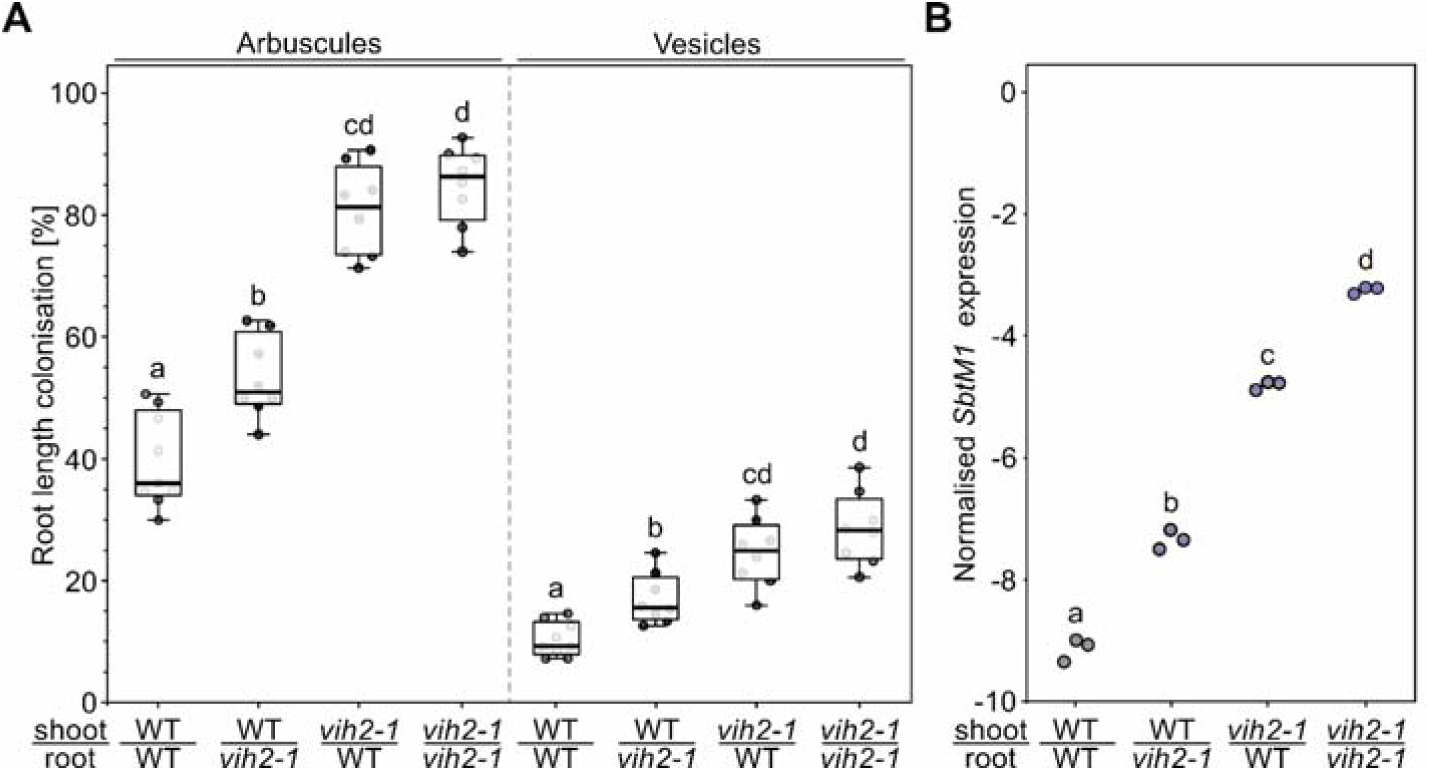
Colonization of *Lotus japonicus* by AM fungi is controlled systemically and locally by (PP)-InsPs. Six-day-old *L. japonicus* wildtype (WT) and *vih2-1* mutant seedlings were self-grafted or reciprocally grafted an grown for three weeks on plates. Subsequently, successful grafts were planted in open pots containing 300 mL washed sand (5 seedlings per pot) and grown in the presence of *R. irregularis* spore inoculum (Symplanta, Germany; 500 spores per plant). Plants were fertilized once a week with liquid Lotus cultivation medium containin 25 µM P_i_ and harvested five weeks after planting. **(A)** Roots were stained with ink-vinegar and AM colonization was quantified. n = 8–9. For statistical analysis, an ordinary one-way ANOVA with Tukey’s multiple comparisons test was performed. Different letters indicate significant differences. **(B)** The expression level of the AM marker gene *SbtM1* was analyzed *via* qRT-PCR. n = 3. For statistical analysis, an ordinary one-way ANOVA with Tukey’s multiple comparisons test was performed. Different letters indicate significant differences.

**Table S1.**
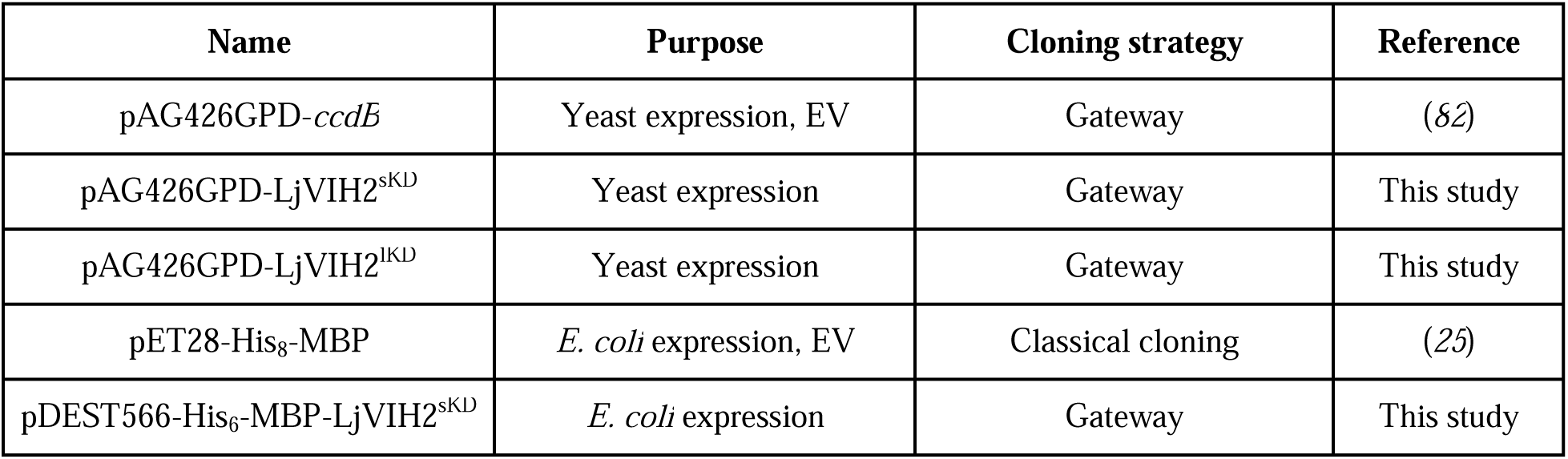
Constructs. Detailed description of the vector backbones and constructs used in the present study produced by Gateway cloning. EV, empty vector control.

| Name | Purpose | Cloning strategy | Reference |
| --- | --- | --- | --- |
| pAG426GPD-*ccdB* | Yeast expression, EV | Gateway | (82) |
| pAG426GPD-LjVIH2<sup>sKD</sup> | Yeast expression | Gateway | This study |
| pAG426GPD-LjVIH2<sup>IKD</sup> | Yeast expression | Gateway | This study |
| pET28-His<sub>8</sub>-MBP | *E. coli* expression, EV | Classical cloning | (25) |
| pDEST566-His<sub>6</sub>-MBP-LjVIH2<sup>sKD</sup> | *E. coli* expression | Gateway | This study |

**Table S2.**
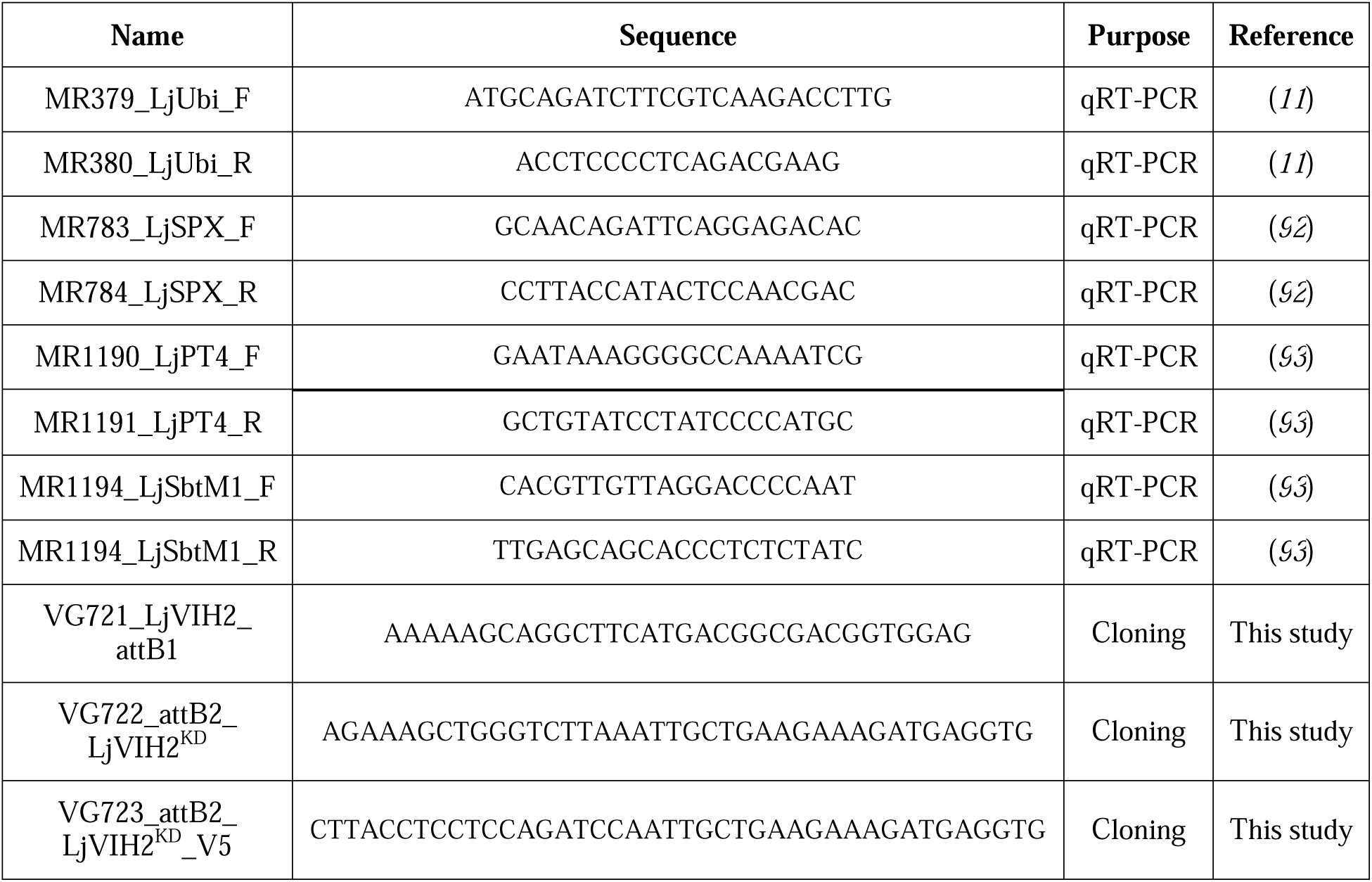
Primers. List of primers used in the present study.

## References and Notes

1. J. Paz-Ares, M. I. Puga, M. Rojas-Triana, I. Martinez-Hevia, S. Diaz, C. Poza-Carrión, M. Miñambres, A. Leyva, Plant adaptation to low phosphorus availability: Core signaling, crosstalks, and applied implications. Mol. Plant 15, 104–124 (2022).

2. C. L. Dybas, Dead Zones Spreading in World Oceans. Bioscience 55, 552–557 (2005).

3. V. Rubio, F. Linhares, R. Solano, A. C. Martín, J. Iglesias, A. Leyva, J. Paz-Ares, A conserved MYB transcription factor involved in phosphate starvation signaling both in vascular plants and in unicellular algae. Genes Dev. 15, 2122–2133 (2001).

4. R. Bustos, G. Castrillo, F. Linhares, M. I. Puga, V. Rubio, J. Pérez-Pérez, R. Solano, A. Leyva, J. Paz-Ares, A central regulatory system largely controls transcriptional activation and repression responses to phosphate starvation in Arabidopsis. PLoS Genet. 6, e1001102 (2010).

5. L. Sun, L. Song, Y. Zhang, Z. Zheng, D. Liu, Arabidopsis PHL2 and PHR1 act redundantly as the key components of the central regulatory system controlling transcriptional responses to phosphate starvation. Plant Physiol. 170, 499–514 (2016).

6. Z. Wang, Z. Zheng, L. Song, D. Liu, Functional Characterization of Arabidopsis PHL4 in plant response to phosphate starvation. Front. Plant Sci. 9, 1432 (2018).

7. F. Ren, Q.-Q. Guo, L.-L. Chang, L. Chen, C.-Z. Zhao, H. Zhong, X.-B. Li, *Brassica napus PHR1* gene encoding a MYB-like protein functions in response to phosphate starvation. PLoS One 7, e44005 (2012).

8. J. Zhou, F. Jiao, Z. Wu, Y. Li, X. Wang, X. He, W. Zhong, P. Wu, *OsPHR2* is involved in phosphate-starvation signaling and excessive phosphate accumulation in shoots of plants. Plant Physiol. 146, 1673–1686 (2008).

9. J. Wang, J. Sun, J. Miao, J. Guo, Z. Shi, M. He, Y. Chen, X. Zhao, B. Li, F. Han, Y. Tong, Z. Li, A phosphate starvation response regulator *Ta-PHR1* is involved in phosphate signalling and increases grain yield in wheat. Ann. Bot. 111, 1139–1153 (2013).

10. P. Wang, R. Snijders, W. Kohlen, J. Liu, T. Bisseling, E. Limpens, Medicago SPX1 and SPX3 regulate phosphate homeostasis, mycorrhizal colonization, and arbuscule degradation. Plant Cell 33, 3470–3486 (2021).

11. D. Das, M. Paries, K. Hobecker, M. Gigl, C. Dawid, H.-M. Lam, J. Zhang, M. Chen, C. Gutjahr, PHOSPHATE STARVATION RESPONSE transcription factors enable arbuscular mycorrhiza symbiosis. Nat. Commun. 13, 477 (2022).

12. Q. Lv, Y. Zhong, Y. Wang, Z. Wang, L. Zhang, J. Shi, Z. Wu, Y. Liu, C. Mao, K. Yi, P. Wu, SPX4 negatively regulates phosphate signaling and homeostasis through its interaction with PHR2 in rice. Plant Cell 26, 1586–1597 (2014).

13. M. I. Puga, I. Mateos, R. Charukesi, Z. Wang, J. M. Franco-Zorrilla, L. de Lorenzo, M. L. Irigoyen, S. Masiero, R. Bustos, J. Rodríguez, A. Leyva, V. Rubio, H. Sommer, J. Paz-Ares, SPX1 is a phosphate-dependent inhibitor of Phosphate Starvation Response 1 in Arabidopsis. Proc. Natl. Acad. Sci. U. S. A. 111, 14947–14952 (2014).

14. Z. Wang, W. Ruan, J. Shi, L. Zhang, D. Xiang, C. Yang, C. Li, Z. Wu, Y. Liu, Y. Yu, H. Shou, X. Mo, C. Mao, P. Wu, Rice SPX1 and SPX2 inhibit phosphate starvation responses through interacting with PHR2 in a phosphate-dependent manner. Proc. Natl. Acad. Sci. U.S.A. 111, 14953–14958 (2014).

15. W. Qi, I. W. Manfield, S. P. Muench, A. Baker, AtSPX1 affects the AtPHR1–DNA-binding equilibrium by binding monomeric AtPHR1 in solution. Biochem. J 474, 3675–3687 (2017).

16. Y. Zhong, Y. Wang, J. Guo, X. Zhu, J. Shi, Q. He, Y. Liu, Y. Wu, L. Zhang, Q. Lv, C. Mao, Rice SPX6 negatively regulates the phosphate starvation response through suppression of the transcription factor PHR2. New Phytol. 219, 135–148 (2018).

17. M. B. Osorio, S. Ng, O. Berkowitz, I. De Clercq, C. Mao, H. Shou, J. Whelan, R. Jost, SPX4 acts on PHR1-dependent and -independent regulation of shoot phosphorus status in Arabidopsis. Plant Physiol. 181, 332–352 (2019).

18. R. Wild, R. Gerasimaite, J.-Y. Jung, V. Truffault, I. Pavlovic, A. Schmidt, A. Saiardi, H. J. Jessen, Y. Poirier, M. Hothorn, A. Mayer, Control of eukaryotic phosphate homeostasis by inositol polyphosphate sensor domains. Science 352, 986–990 (2016).

19. M. K. Ried, R. Wild, J. Zhu, J. Pipercevic, K. Sturm, L. Broger, R. K. Harmel, L. A. Abriata, L. A. Hothorn, D. Fiedler, S. Hiller, M. Hothorn, Inositol pyrophosphates promote the interaction of SPX domains with the coiled-coil motif of PHR transcription factors to regulate plant phosphate homeostasis. Nat. Commun. 12, 384 (2021).

20. J. Dong, G. Ma, L. Sui, M. Wei, V. Satheesh, R. Zhang, S. Ge, J. Li, T. E. Zhang, C. Wittwer, H. J. Jessen, H. Zhang, G. Y. An, D. Y. Chao, D. Liu, M. Lei, Inositol Pyrophosphate InsP_8_ acts as an intracellular phosphate signal in Arabidopsis. Mol. Plant 12, 1463–1473 (2019).

21. V. Raboy, *Myo*-inositol-1,2,3,4,5,6-hexakisphosphate. Phytochemistry 64, 1033–1043 (2003).

22. S. B. Shears, Inositol pyrophosphates: why so many phosphates? Adv. Biol. Regul. 57, 203–216 (2015).

23. J. Zhu, K. Lau, R. Puschmann, R. K. Harmel, Y. Zhang, V. Pries, P. Gaugler, L. Broger, A. K. Dutta, H. J. Jessen, G. Schaaf, A. R. Fernie, L. A. Hothorn, D. Fiedler, M. Hothorn, Two bifunctional inositol pyrophosphate kinases/phosphatases control plant phosphate homeostasis. Elife 8 (2019).

24. E. Riemer, D. Qiu, D. Laha, R. K. Harmel, P. Gaugler, V. Gaugler, M. Frei, M.-R. Hajirezaei, N. P. Laha, L. Krusenbaum, R. Schneider, A. Saiardi, D. Fiedler, H. J. Jessen, G. Schaaf, R. F. H. Giehl, ITPK1 is an InsP6/ADP phosphotransferase that controls phosphate signaling in Arabidopsis. Mol. Plant 14, 1864–1880 (2021).

25. D. Laha, P. Johnen, C. Azevedo, M. Dynowski, M. Weiß, S. Capolicchio, H. Mao, T. Iven, M. Steenbergen, M. Freyer, P. Gaugler, M. K. F. de Campos, N. Zheng, I. Feussner, H. J. Jessen, S. C. M. Van Wees, A. Saiardi, G. Schaaf, VIH2 Regulates the synthesis of inositol pyrophosphate InsP_8_ and jasmonate-dependent defenses in Arabidopsis. Plant Cell 27, 1082–1097 (2015).

26. H. Wang, V. S. Nair, A. A. Holland, S. Capolicchio, H. J. Jessen, M. K. Johnson, S. B. Shears, Asp1 from *Schizosaccharomyces pombe* binds a [2Fe-2S]^2+^ cluster which inhibits inositol pyrophosphate 1-phosphatase activity. Biochemistry 54, 6462–6474 (2015).

27. M. Pascual-Ortiz, A. Saiardi, E. Walla, V. Jakopec, N. A. Künzel, I. Span, A. Vangala, U. Fleig, Asp1 bifunctional activity modulates spindle function *via* controlling cellular inositol pyrophosphate levels in *Schizosaccharomyces pombe*. Mol. Cell. Biol. 38, 1–21 (2018).

28. D. E. Dollins, W. Bai, P. C. Fridy, J. C. Otto, J. L. Neubauer, S. G. Gattis, K. P. M. Mehta, J. D. York, Vip1 is a kinase and pyrophosphatase switch that regulates inositol diphosphate signaling. Proc. Natl. Acad. Sci. U.S.A. 117, 9356–9364 (2020).

29. E. Riemer, N. J. Pullagurla, R. Yadav, P. Rana, H. J. Jessen, M. Kamleitner, G. Schaaf, D. Laha, Regulation of plant biotic interactions and abiotic stress responses by inositol polyphosphates. Front. Plant Sci. 13, 944515 (2022).

30. S. Smith, D. Read, Mycorrhizal Symbiosis. (Academic Press, 2008).

31. M. C. Brundrett, L. Tedersoo, Evolutionary history of mycorrhizal symbioses and global host plant diversity. New Phytol. 220, 1108–1115 (2018).

32. W. Remy, T. N. Taylor, H. Hass, H. Kerp, Four hundred-million-year-old vesicular arbuscular mycorrhizae. Proc. Natl. Acad. Sci. U.S.A. 91, 11841–11843 (1994).

33. B. J. W. Mills, S. A. Batterman, K. J. Field, Nutrient acquisition by symbiotic fungi governs Palaeozoic climate transition. Philos. Trans. R. Soc. Lond. B Biol. Sci. 373 (2018).

34. A. Genre, M. Chabaud, T. Timmers, P. Bonfante, D. G. Barker, Arbuscular mycorrhizal fungi elicit a novel intracellular apparatus in *Medicago truncatula* root epidermal cells before infection. Plant Cell 17, 3489–3499 (2005).

35. A. Genre, M. Chabaud, A. Faccio, D. G. Barker, P. Bonfante, Prepenetration apparatus assembly precedes and predicts the colonization patterns of arbuscular mycorrhizal fungi within the root cortex of both *Medicago truncatula* and *Daucus carota*. Plant Cell 20, 1407–1420 (2008).

36. M. J. Harrison, Cellular programs for arbuscular mycorrhizal symbiosis. Curr. Opin. Plant Biol. 15, 691–698 (2012).

37. C. Gutjahr, M. Parniske, Cell and developmental biology of arbuscular mycorrhiza symbiosis. Annu. Rev. Cell Dev. Biol. 29, 593–617 (2013).

38. S. Campo, B. San Segundo, Systemic induction of phosphatidylinositol-based signaling in leaves of arbuscular mycorrhizal rice plants. Sci. Rep. 10, 15896 (2020).

39. D. Liao, C. Sun, H. Liang, Y. Wang, X. Bian, C. Dong, X. Niu, M. Yang, G. Xu, A. Chen, S. Wu, SlSPX1-SlPHR complexes mediate the suppression of arbuscular mycorrhizal symbiosis by phosphate repletion in tomato. Plant Cell 34, 4045–4065 (2022).

40. J. Shi, B. Zhao, S. Zheng, X. Zhang, X. Wang, W. Dong, Q. Xie, G. Wang, Y. Xiao, F. Chen, N. Yu, E. Wang, A phosphate starvation response-centered network regulates mycorrhizal symbiosis. Cell 184, 5527–5540.e18 (2021).

41. P. Wang, Y. Zhong, Y. Li, W. Zhu, Y. Zhang, J. Li, Z. Chen, E. Limpens, The phosphate starvation response regulator PHR2 antagonizes arbuscule maintenance in Medicago. New Phytol., doi: 10.1111/nph.19869 (2024).

42. F. Lota, S. Wegmüller, B. Buer, S. Sato, A. Bräutigam, B. Hanf, M. Bucher, The *cis*-acting CTTC-P1BS module is indicative for gene function of *LjVTI12*, a Qb-SNARE protein gene that is required for arbuscule formation in *Lotus japonicus*. Plant J. 74, 280–293 (2013).

43. A. Chen, M. Gu, S. Sun, L. Zhu, S. Hong, G. Xu, Identification of two conserved *cis*-acting elements, MYCS and P1BS, involved in the regulation of mycorrhiza-activated phosphate transporters in eudicot species. New Phytol. 189, 1157–1169 (2011).

44. A. Bravo, M. Brands, V. Wewer, P. Dörmann, M. J. Harrison, Arbuscular mycorrhiza-specific enzymes FatM and RAM2 fine-tune lipid biosynthesis to promote development of arbuscular mycorrhiza. New Phytol. 214, 1631–1645 (2017).

45. L. H. Luginbuehl, G. N. Menard, S. Kurup, H. Van Erp, G. V. Radhakrishnan, A. Breakspear, G. E. D. Oldroyd, P. J. Eastmond, Fatty acids in arbuscular mycorrhizal fungi are synthesized by the host plant. Science 356, 1175–1178 (2017).

46. Y. Jiang, W. Wang, Q. Xie, N. Liu, L. Liu, D. Wang, X. Zhang, C. Yang, X. Chen, D. Tang, E. Wang, Plants transfer lipids to sustain colonization by mutualistic mycorrhizal and parasitic fungi. Science 356, 1172–1175 (2017).

47. R. Roth, U. Paszkowski, Plant carbon nourishment of arbuscular mycorrhizal fungi. Curr. Opin. Plant Biol. 39, 50–56 (2017).

48. A. Keymer, P. Pimprikar, V. Wewer, C. Huber, M. Brands, S. L. Bucerius, P.-M. Delaux, V. Klingl, E. von Röpenack-Lahaye, T. L. Wang, W. Eisenreich, P. Dörmann, M. Parniske, C. Gutjahr, Lipid transfer from plants to arbuscular mycorrhiza fungi. Elife 6 (2017).

49. S. Ivanov, M. J. Harrison, Receptor-associated kinases control the lipid provisioning program in plant-fungal symbiosis. Science. 383, 443–448 (2024).

50. S. E. Smith, F. A. Smith, Roles of arbuscular mycorrhizas in plant nutrition and growth: new paradigms from cellular to ecosystem scales. Annu. Rev. Plant Biol. 62, 227–250 (2011).

51. N. Marro, G. Grilli, F. Soteras, M. Caccia, S. Longo, N. Cofré, V. Borda, M. Burni, M. Janoušková, C. Urcelay, The effects of arbuscular mycorrhizal fungal species and taxonomic groups on stressed and unstressed plants: a global meta-analysis. New Phytol. 235, 320–332 (2022).

52. F. Breuillin, J. Schramm, M. Hajirezaei, A. Ahkami, P. Favre, U. Druege, B. Hause, M. Bucher, T. Kretzschmar, E. Bossolini, C. Kuhlemeier, E. Martinoia, P. Franken, U. Scholz, D. Reinhardt, Phosphate systemically inhibits development of arbuscular mycorrhiza in *Petunia hybrida* and represses genes involved in mycorrhizal functioning. Plant J. 64, 1002–1017 (2010).

53. C. Balzergue, V. Puech-Pagès, G. Bécard, S. F. Rochange, The regulation of arbuscular mycorrhizal symbiosis by phosphate in pea involves early and systemic signalling events. J. Exp. Bot. 62, 1049–1060 (2011).

54. S. Osada, K. Kageyama, Y. Ohnishi, J.-I. Nishikawa, T. Nishihara, M. Imagawa, Inositol phosphate kinase Vip1p interacts with histone chaperone Asf1p in *Saccharomyces cerevisiae*. Mol. Biol. Rep. 39, 4989–4996 (2012).

55. A. S. Mulugu, W. Bai, P. C. Fridy, R. J. Bastidas, J. C. Otto, D. E. Dollins, T. A. Haystead, A. Ribeiro, J. D. York, A conserved family of enzymes that phosphorylate inositol hexakisphosphate. Science 316, 106–109 (2007).

55. S. M. N. Onnebo, A. Saiardi, Inositol pyrophosphates modulate hydrogen peroxide signalling. Biochem. J. 423, 109–118 (2009).

56. G. Liu, E. Riemer, R. Schneider, D. Cabuzu, O. Bonny, C. A. Wagner, D. Qiu, A. Saiardi, A. Strauss, T. Lahaye, G. Schaaf, T. Knoll, J. P. Jessen, H. J. Jessen, The phytase RipBL1 enables the assignment of a specific inositol phosphate isomer as a structural component of human kidney stones. *RSC Chem*. Biol. 4, 300–309 (2023).

57. D. Qiu, C. Gu, G. Liu, K. Ritter, V. B. Eisenbeis, T. Bittner, A. Gruzdev, L. Seidel, B. Bengsch, S. B. Shears, H. J. Jessen, Capillary electrophoresis mass spectrometry identifies new isomers of inositol pyrophosphates in mammalian tissues. Chem. Sci. 14, 658–667 (2023).

58. D. Qiu, V. B. Eisenbeis, A. Saiardi, H. J. Jessen, Absolute quantitation of inositol pyrophosphates by capillary electrophoresis electrospray ionization mass spectrometry. J. Vis. Exp., doi: 10.3791/62847-v (2021).

59. D. Qiu, M. S. Wilson, V. B. Eisenbeis, R. K. Harmel, E. Riemer, T. M. Haas, C. Wittwer, N. Jork, C. Gu, S. B. Shears, G. Schaaf, B. Kammerer, D. Fiedler, A. Saiardi, H. J. Jessen, Analysis of inositol phosphate metabolism by capillary electrophoresis electrospray ionization mass spectrometry. Nat. Commun. 11, 6035 (2020).

60. A. Małolepszy, T. Mun, N. Sandal, V. Gupta, M. Dubin, D. Urbański, N. Shah, A. Bachmann, E. Fukai, H. Hirakawa, S. Tabata, M. Nadzieja, K. Markmann, J. Su, Y. Umehara, T. Soyano, A. Miyahara, S. Sato, M. Hayashi, J. Stougaard, S. U. Andersen, The LORE1 insertion mutant resource. Plant J. 88, 306–317 (2016).

61. T. Mun, A. Bachmann, V. Gupta, J. Stougaard, S. U. Andersen, Lotus Base: An integrated information portal for the model legume *Lotus japonicus*. Sci. Rep. 6, 39447 (2016).

62. P. Gaugler, V. Gaugler, M. Kamleitner, G. Schaaf, Extraction and quantification of soluble, radiolabeled inositol polyphosphates from different plant species using SAX-HPLC. J. Vis. Exp., doi: 10.3791/61495 (2020).

63. L. Arata, E. Fabrizi, P. Sckokai, A worldwide analysis of trend in crop yields and yield variability: Evidence from FAO data. Econ. Model. 90, 190–208 (2020).

64. R. J. H. Sawers, C. Gutjahr, U. Paszkowski, Cereal mycorrhiza: an ancient symbiosis in modern agriculture. Trends Plant Sci. 13, 93–97 (2008).

65. E. Verbruggen, E. Toby Kiers, Evolutionary ecology of mycorrhizal functional diversity in agricultural systems. Evol. Appl. 3, 547–560 (2010).

66. M. Moora, J. Davison, M. Öpik, M. Metsis, Ü. Saks, T. Jairus, M. Vasar, M. Zobel, Anthropogenic land use shapes the composition and phylogenetic structure of soil arbuscular mycorrhizal fungal communities. FEMS Microbiol. Ecol. 90, 609–621 (2014).

67. D. Xiang, E. Verbruggen, Y. Hu, S. D. Veresoglou, M. C. Rillig, W. Zhou, T. Xu, H. Li, Z. Hao, Y. Chen, B. Chen, Land use influences arbuscular mycorrhizal fungal communities in the farming-pastoral ecotone of northern China. New Phytol. 204, 968–978 (2014).

68. S. M. Schmidt, M. Belisle, W. B. Frommer, The evolving landscape around genome editing in agriculture: Many countries have exempted or move to exempt forms of genome editing from GMO regulation of crop plants: Many countries have exempted or move to exempt forms of genome editing from GMO regulation of crop plants. EMBO Rep. 21, e50680 (2020).

69. N. P. Laha, R. F. H. Giehl, E. Riemer, D. Qiu, N. J. Pullagurla, R. Schneider, Y. W. Dhir, R. Yadav, Y. E. Mihiret, P. Gaugler, V. Gaugler, H. Mao, N. Zheng, N. von Wirén, A. Saiardi, S. Bhattacharjee, H. J. Jessen, D. Laha, G. Schaaf, INOSITOL (1,3,4) TRIPHOSPHATE 5/6 KINASE1-dependent inositol polyphosphates regulate auxin responses in Arabidopsis. Plant Physiol. 190, 2722–2738 (2022).

70. P. Gaugler, R. Schneider, G. Liu, D. Qiu, J. Weber, J. Schmid, N. Jork, M. Häner, K. Ritter, N. Fernández-Rebollo, R. F. H. Giehl, M. N. Trung, R. Yadav, D. Fiedler, V. Gaugler, H. J. Jessen, G. Schaaf, D. Laha, Arabidopsis PFA-DSP-type phosphohydrolases target specific inositol pyrophosphate messengers. Biochemistry 61, 1213–1227 (2022).

71. F. Laurent, S. M. Bartsch, A. Shukla, F. Rico-Resendiz, D. Couto, C. Fuchs, J. Nicolet, S. Loubéry, H. J. Jessen, D. Fiedler, M. Hothorn, Inositol pyrophosphate catabolism by three families of phosphatases regulates plant growth and development. PLoS Genet. 20, e1011468 (2024).

72. M. Bigalke, A. Ulrich, A. Rehmus, A. Keller, Accumulation of cadmium and uranium in arable soils in Switzerland. Environ. Pollut. 221, 85–93 (2017).

73. G. A. Khan, S. Bouraine, S. Wege, Y. Li, M. de Carbonnel, P. Berthomieu, Y. Poirier, H. Rouached, Coordination between zinc and phosphate homeostasis involves the transcription factor PHR1, the phosphate exporter PHO1, and its homologue PHO1;H3 in Arabidopsis. J. Exp. Bot. 65, 871–884 (2014).

74. W. Zhang, W. Zhang, X. Wang, D. Liu, C. Zou, X. Chen, Quantitative evaluation of the grain zinc in cereal crops caused by phosphorus fertilization. A meta-analysis. Agron. Sustain. Dev. 41 (2021).

75. E. A. Ova, U. B. Kutman, L. Ozturk, I. Cakmak, High phosphorus supply reduced zinc concentration of wheat in native soil but not in autoclaved soil or nutrient solution. Plant Soil 393, 147–162 (2015).

76. J. Ding, L. Liu, C. Wang, L. Shi, F. Xu, H. Cai, High level of zinc triggers phosphorus starvation by inhibiting root-to-shoot translocation and preferential distribution of phosphorus in rice plants. Environ. Pollut. 277, 116778 (2021).

77. F. T. Bingham, M. J. Garber, Solubility and availability of micronutrients in relation to phosphorus fertilization. Soil Sci. Soc. Am. J. 24, 209–213 (1960).

78. B. Yu, C. Zhou, Z. Wang, M. Bucher, G. Schaaf, R. J. H. Sawers, X. Chen, F. Hochholdinger, C. Zou, P. Yu, Maize zinc uptake is influenced by arbuscular mycorrhizal symbiosis under various soil phosphorus availabilities. New Phytol. 243, 1936–1950 (2024).

79. M. Abdalla, M. Bitterlich, J. Jansa, D. Püschel, M. A. Ahmed, The role of arbuscular mycorrhizal symbiosis in improving plant water status under drought. J. Exp. Bot. 74, 4808–4824 (2023).

80. H.-J. Hawkins, R. I. M. Cargill, M. E. Van Nuland, S. C. Hagen, K. J. Field, M. Sheldrake, N. A. Soudzilovskaia, E. T. Kiers, Mycorrhizal mycelium as a global carbon pool. Curr. Biol. 33, R560–R573 (2023).

81. S. Alberti, A. D. Gitler, S. Lindquist, A suite of Gateway cloning vectors for high-throughput genetic analysis in *Saccharomyces cerevisiae*. Yeast 24, 913–919 (2007).

82. H. Vierheilig, A. P. Coughlan, U. Wyss, Y. Piché, Ink and vinegar, a simple staining technique for arbuscular-mycorrhizal fungi. Appl. Environ. Microbiol. 64, 5004–5007 (1998).

83. T. P. McGonigle, M. H. Miller, D. G. Evans, G. L. Fairchild, J. A. Swan, A new method which gives an objective measure of colonization of roots by vesicular-arbuscular mycorrhizal fungi. New Phytol. 115, 495–501 (1990).

84. D. Dreher, H. Yadav, S. Zander, B. Hause, Is there genetic variation in mycorrhization of *Medicago truncatula*? PeerJ 5, e3713 (2017).

85. M. Sexauer, H. Bhasin, M. Schön, E. Roitsch, C. Wall, U. Herzog, K. Markmann, A micro RNA mediates shoot control of root branching. Nat. Commun. 14, 8083 (2023).

86. R. D. Gietz, R. H. Schiestl, A. R. Willems, R. A. Woods, Studies on the transformation of intact yeast cells by the LiAc/SS-DNA/PEG procedure. Yeast 11, 355–360 (1995).

87. B. J. M. Zonneveld, Cheap and simple yeast media. J. Microbiol. Methods 4, 287–291 (1986).

88. C. Azevedo, A. Saiardi, Extraction and analysis of soluble inositol polyphosphates from yeast. Nat. Protoc. 1, 2416–2422 (2006).

89. O. Losito, Z. Szijgyarto, A. C. Resnick, A. Saiardi, Inositol pyrophosphates and their unique metabolic complexity: analysis by gel electrophoresis. PLoS One 4, e5580 (2009).

90. R. Schneider, K. Lami, I. Prucker, S. C. Stolze, A. Strauß, K. Langenbach, M. Kamleitner, Y. Z. Belay, K. Ritter, D. Furkert, P. Gaugler, E. Lange, N. Faiß, J. M. Schmidt, M. Harings, L. Krusenbaum, S. Wege, S. Kriescher, J. The, H. Schoof, D. Fiedler, H. Nakagami, R. F. H. Giehl, T. Lahaye, H. J. Jessen, V. Gaugler, G. Schaaf, NUDIX hydrolases target specific inositol pyrophosphates and regulate phosphate and iron homeostasis, and the expression of defense genes in Arabidopsis, bioRxiv (2024). doi: 10.1101/2024.10.18.619122.

91. V. Volpe, M. Giovannetti, X.-G. Sun, V. Fiorilli, P. Bonfante, The phosphate transporters LjPT4 and MtPT4 mediate early root responses to phosphate status in non-mycorrhizal roots. Plant Cell Environ. 39, 660–671 (2016).

92. M. Groth, S. Kosuta, C. Gutjahr, K. Haage, S. L. Hardel, M. Schaub, A. Brachmann, S. Sato, S. Tabata, K. Findlay, T. L. Wang, M. Parniske, Two *Lotus japonicus* symbiosis mutants impaired at distinct steps of arbuscule development. Plant J. 75, 117–129 (2013).

